# Evaluation of Alphafold modeling for elucidation of nanobody-peptide epitope interactions

**DOI:** 10.1101/2025.03.14.643405

**Authors:** Shivani Sachdev, Swarnali Roy, Shubhra Saha, Gengxiang Zhao, Rashmi Kumariya, Brendan A. Creemer, Rui Yin, Brian G. Pierce, Carole A. Bewley, Ross W. Cheloha

## Abstract

Models of Ab-antigen complexes can be used to understand interaction mechanisms and for improving affinity. This study evaluates the use of the protein structure prediction algorithm AlphaFold (AF) for exploration of interactions between peptide epitope tags and the smallest functional antibody fragments, nanobodies (Nbs). Although past studies of AF for modeling antibody-target (antigen) interactions suggested modest algorithm performance, those were primarily focused on Ab-protein interactions, while the performance and utility of AF for Nb-peptide interactions, which are generally less complex due to smaller antigens, smaller binding domains, and fewer chains, is less clear. In this study we evaluated the performance of AF for predicting the structures of Nbs bound to experimentally validated, linear, short peptide epitopes (Nb-tag pairs). We expanded the pool of experimental data available for comparison through crystallization and structural determination of a previously reported Nb-tag complex (Nb_127_). Models of Nb-tag pair structures generated from AF were variable with respect to consistency with experimental data, with good performance in just over half (4 out of 6) of cases. Even among Nb-tag pairs successfully modeled in isolation, efforts to translate modeling to more complex contexts failed, suggesting an underappreciated role of the size and complexity of inputs in AF modeling success. Finally, the model of a Nb-tag pair with minimal previous characterization was used to guide the design of a peptide-electrophile conjugate that undergoes covalent crosslinking with Nb upon binding. These findings highlight the utility of minimized antibody and antigen structures to maximize insights from AF modeling.

## Introduction

Decades of intensive study have provided a rich collection of structural data that link protein primary structure (sequence) with structure.^1^ In parallel, impressive advances in technologies for DNA sequencing have generated a compendium of sequence information covering a large swath of organisms alive today.^2^ Modern computational methodology has leveraged rich libraries of DNA sequence information, evolutionary analysis, and structural databases to generate algorithms for predicting structures of previously unanalyzed or fully novel proteins.^3,4^ Protein structure prediction methodology has realized substantial improvements in speed and performance following the publication and widespread implementation of the AlphaFold 2 (AF2) algorithm.^5^ Refinements of this methodology have enabled high quality predictions of the structures of proteins involved in protein-protein interactions (PPIs).^6^ One important class of PPIs that has often been poorly modeled via computation is the interaction between antibodies (Ab) and their targets (antigens).^7^ Notably, AF2 performed better than other comparable modeling methods, although there remains room for improvement.^8,9^ Further improvement in these efforts has been realized with the release of AlphaFold3 (AF3).^10^ This work focuses on application of AF2 given its widespread characterization and the abundance of tools available for easy implementation.

Ab-antigen interactions represent a challenge for prediction algorithms that use co-evolution of amino acid residues for predicting protein folding and PPI interfaces, such as AF2.^11^ Genetic recombination, somatic (hyper)mutation, and clonal selection drive sequence diversification of Abs produced by the immune system over a timeframe that is short by human evolutionary standards. For example, a single immunization or infection in humans can result in generation of many novel Ab sequences that bind to antigens through new or unusual mechanisms.^12^ This complex process of immune system-mediated Ab sequence variation, typically without a change in antigen sequence, and its impact on binding to antigens of interest is incompletely captured in sequence data used to train modeling algorithms. Further difficulty in modeling arises from the mechanism through which Abs bind to their targets. The binding surface for Abs occurs primarily over six complementarity determining regions (CDRs), loops that are evenly distributed between the Ab heavy chain (HC) and light chain (LC) (**Figure 1A**).^13^ Simplified systems to assess modelling Ab-antigen interactions could be useful for guiding improvements in modeling approaches.

**Figure 1.**
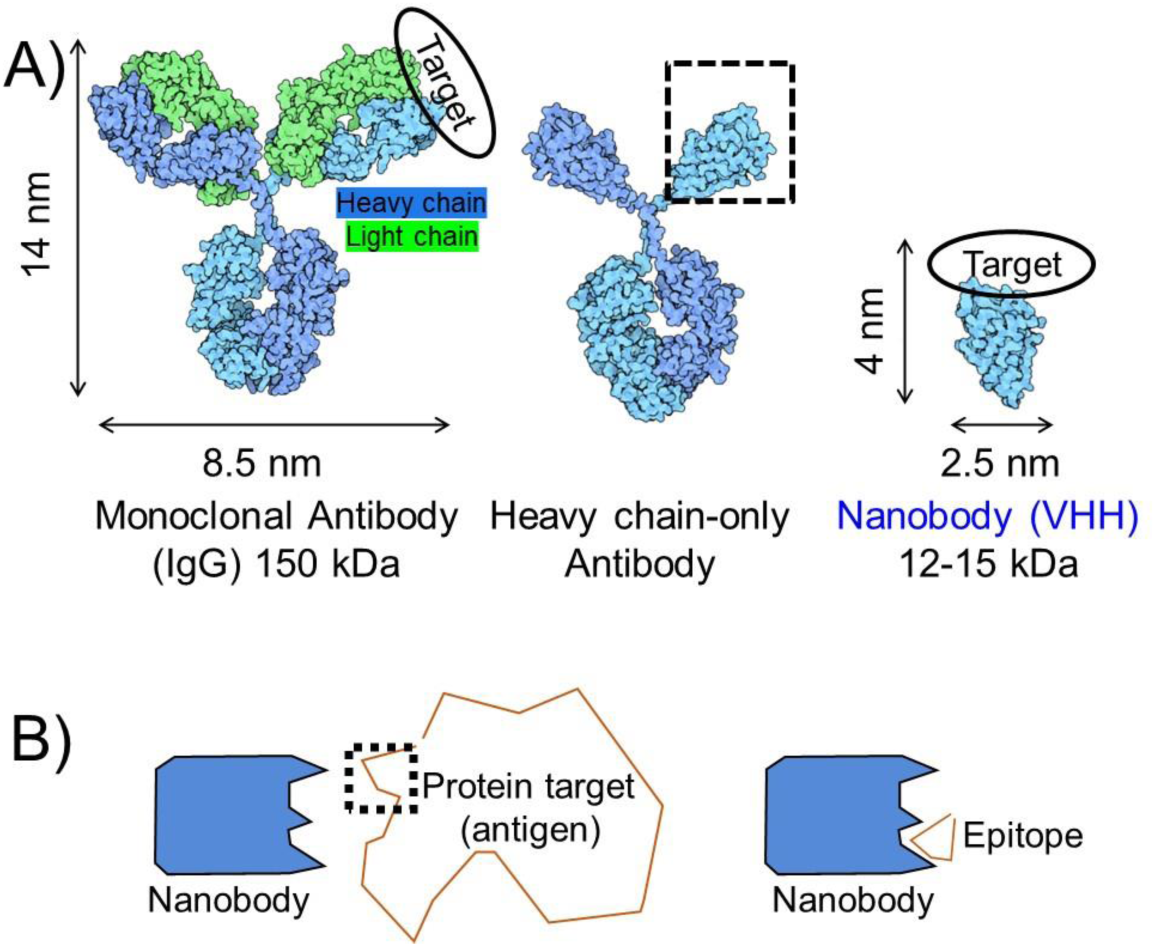
Illustration of antibody and antigen complexity. **A)** Comparison of full-sized antibodies to heavy chain-only antibodies and nanobodies (Nbs). **B)** Comparison of full-sized protein antigens to epitope fragments.

Unlike conventional Abs from humans and mice (IgGs), which rely on an Ab structure comprised of HC and LC polypeptides, camelids (camels, alpacas, llamas) also produce a type of Ab made only from HC.^14^ Although the physiological role of HC-only Abs is not clear, the single chain architecture has proven valuable for many biological applications.^15^ Excision of the antigen binding domain of HC-only antibodies provides variable heavy chain domains of HC-only Abs (VHHs, also known as nanobodies, **Figure 1A**). Nanobodies (Nbs) represent the smallest Ab fragment that maintains high antigen binding affinity. Nbs are 10% the size of conventional IgGs and unlike IgGs they frequently do not require disulfide bonds or glycosylation for folding or function.^16^ Because of these features Nbs can be recombinantly expressed in bacteria in high yield. Nbs have been widely used to stabilize conformationally flexible proteins for structural studies and for imaging applications.^17,18^ Site-specific labeling of Nbs has been achieved using a variety of chemical and enzymatic approaches, which has facilitated the production of Nb-ligand conjugates for the study of cell surface receptor signaling.^19–22^ Features of Nbs potentially useful for modeling applications are their small size, rigid single domain architecture, and binding mechanism that relies on only three CDRs (vs. six for conventional IgGs). Indeed, a higher level of success was observed in modeling of Nb-antigen complexes compared to Ab-antigen complexes.^8^

Antigen size and complexity constitutes another variable that is potentially important to the success of modelling Ab-antigen interactions. Although many Abs and Nbs bind to a discontinuous (conformational) epitope on their target, a substantial fraction bind primarily through a continuous stretch of amino acids (linear epitope). For some linear epitopes, these sequences can be extracted and synthesized chemically or fused to a partner protein for evaluation of binding outside the context of the natural protein antigen. This approach is often used in epitope tagging, wherein target proteins for which high quality antibody detection reagents are unavailable, are modified to contain an epitope tag recognized by a high-quality antibody.^23^ Evaluation of Abs or Nbs that bind short linear epitopes offers the possibility of comparing the modeling output generated for Ab-full size antigen complexes to those of Ab-epitope complexes (**Figure 1b**). In this work, we analyze previously identified Nb-linear peptide epitope (tag) interactions^24–32^ through AF2 modelling. We restrict our analysis to Nb-peptide epitope pairs (six total) that meet one of these criteria: experimental structural information is available, substantial structure-activity relationship studies have been performed, or our group is actively involved in their characterization. By restricting our analysis to Nbs that bind to small peptide epitopes we highlight the important role of epitope complexity in modelling success and the unpredictable nature of modelling epitope recognition in different protein contexts. We further expand this line of inquiry by characterizing for the first time the structure of a Nb-tag complex via X-ray crystallographic analysis. Further, we demonstrate that AF can be used to predict the binding of Nbs to variants of peptide epitopes, which results in the identification of analogues with improved Nb affinity. Finally, we show that AF modeling can be used to guide the design of epitope peptide-electrophile conjugates that form a covalent bond with target upon Nb binding.^33^

## Results

We performed a survey of the literature to compile a collection of Nbs that bind to short (< 20 residue), linear peptide epitopes. We focused on Nb-peptide interactions for which structural information was available (**Figure 2**) or those used previously for in-house studies (**Figure 3, Supporting Table 1**).^24–32^ We compared four structurally characterized nanobody-peptide complexes to structural models generated using AF2 multimer version 3 in ColabFold.^34^ Unless otherwise indicated, all models were generated using this implementation of Colabfold. Of the four experimental Nb-tag structures, all were deposited into online repositories prior to date (2021-09-30) at which the Alphafold2 multimer version 3 training set was extracted (**Supporting Table 1**). We used AF2 to generate an ensemble of five structural models for each of these Nb-peptide tag complexes. We superimposed the ensemble (**Figure 2a**, middle) and the experimental structure (**Figure 2a**, right) using Pymol. Visual inspection shows a substantial variation in consistency for both the ensemble and model-experimental alignments. For the Nb_Alfa_-tag interaction there was a close alignment between all the modelled complexes and the experimentally characterized complex (**Figure 2a**).^28^ Nb_PepTag_ models were more variable, with one model (rank 1, cyan) closely resembling the experimental structure and others showing more divergence (**Figure 2a**). The other Nb-epitope pairs modeled showed major discrepancies with experimental data. For Nb_headlock_ and Nb_MoonTag_ there were notable differences between the experimentally determined complex structures^26^ and the structures generated by modelling (**Figure 2a**). Past work^7,8^ showed that confidence parameters generated by AlphaFold offer predictive power for assessing protein-protein complex model accuracy. We observe an analogous trend here (**Supporting Table 2**). Colabfold models with interface template modeling scores (ipTM) scores above 0.8 (all Nb_Alfa_ models and the top ranked Nb_PepTag_ model) all showed good agreement with experimental structures, with interface and peptide root-mean-square distance (RMSD) values below 2 Å and Medium or High accuracy based on CAPRI criteria.^35^ In contrast, complexes with lower ipTM scores showed lower accuracy, measured by RMSD and by visual inspection of overlayed models (**Figure 2a**).

**Figure 2.**
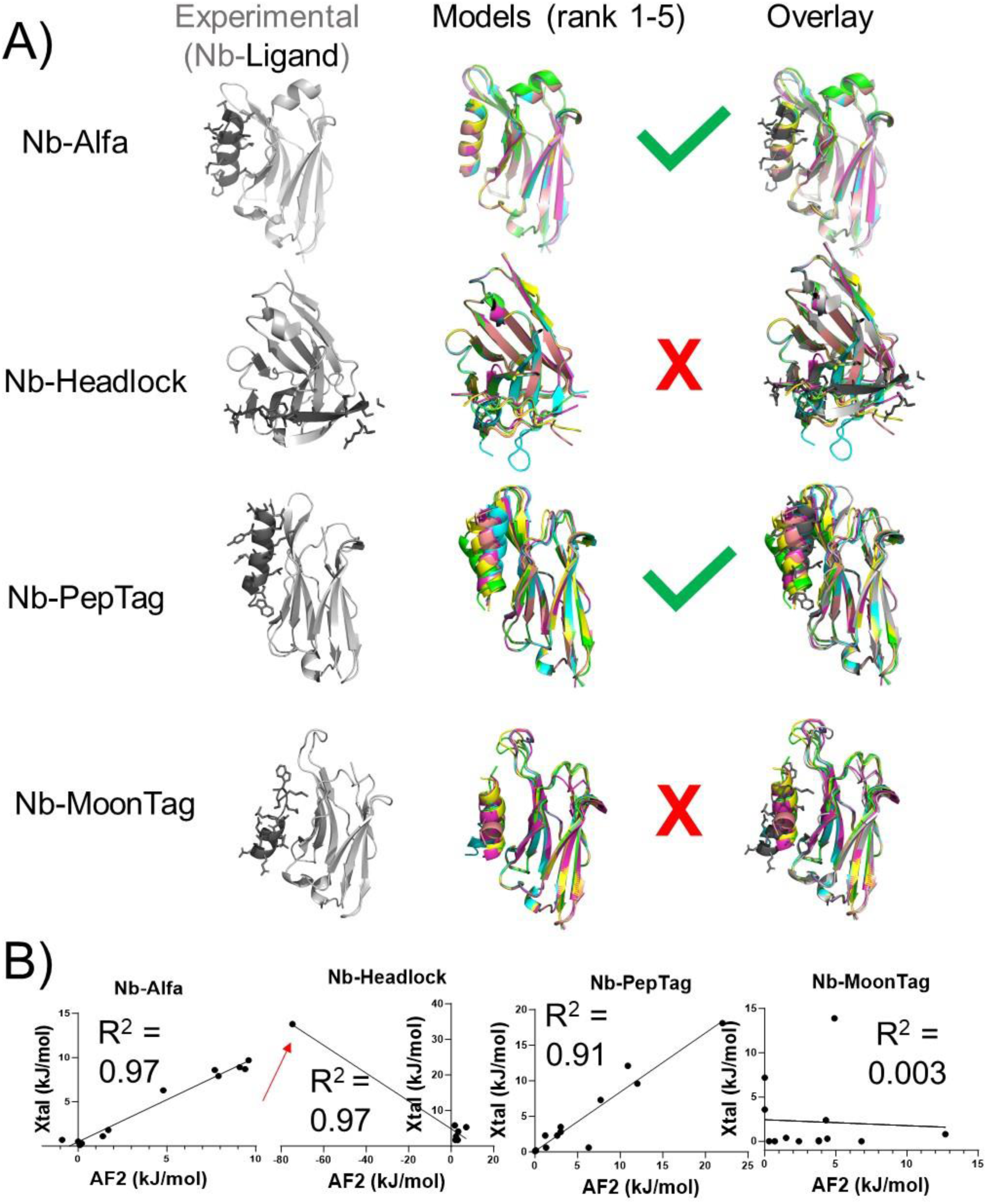
Comparison of experimental Nb-epitope structures to AF2-generated models. **A)** Experimental structures (gray and black, left) are compared to an overlay of 5 complex models produced by AF2 (middle, color) via overlay (right). Structures and overlays were produced using Pymol. **B)** Quantitative assessment of peptide epitope side chain energetic contributions to binding. The energetic contribution of each side chain within experimental or the top ranked AF2 generated complex was quantified using BUDE Ala Scan. Each side chain was plotted as a single point in a scatter plot. A linear correlation model was used to generate a trendline in GraphPad Prism. A red arrow highlights a residues that form a steric clash in a AF2 generated model.

**Figure 3.**
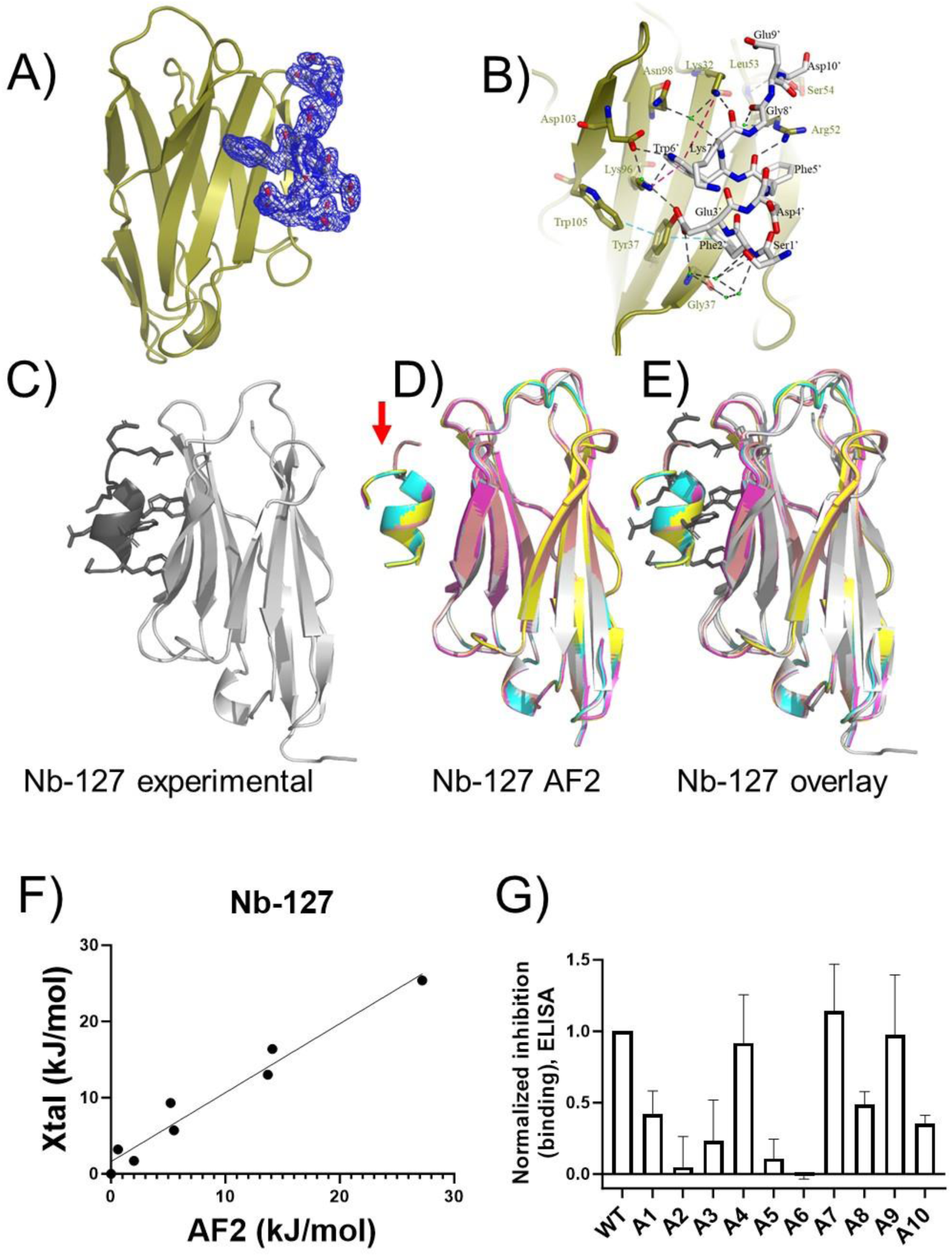
Comparison of a new Nb_127_-eptiope peptide complex structure to AF2-generated models. A complex of Nb_127_ and 127-tag was crystallized and analyzed as described in Methods. **A)** Depiction of peptide epitope electron density. The Nb is shown in cartoon and the 127-tag in sticks with σ=1 map in blue mesh. **B)** Detailed map of Nb_127_-peptide interaction. 127-tag is shown in sticks (black text for residue labels) and Nb in cartoon, with residues of Nb interacting with 127-tag in sticks with gold text labels. Black dashes show hydrogen bonding interaction within the complex. The cyan dashed line shows a pi-pi contact between Trp105, Tyr37 and Phe2’ (4.7 Å and 5.0 Å respectively). The red dashed line shows the Pi-cation contact between Lys32, Lys96 and Trp6’ (3.3 Å and 6.5 Å, respectively). Green balls show waters involved in intermediating the H-bond between Nb and 127-tag. **C-E**) An experimental structure (panel C) is compared to an overlay of 5 complex models produced by AF2 (panel D) via overlay (panel E). **F**) Quantitative assessment of peptide epitope side chain energetic contributions to binding, performed as described in Figure 2 and Methods. **G**) Experimental evaluation of the binding of 127-tag variants to Nb_127_ using ELISA. Peptide characterization is shown in **Supporting Table 5**. Higher bars correspond to more complete inhibition and stronger binding (see Methods). Data correspond to mean ± SD from three independent replicates.

These comparisons, and specific amino acid side chains predicted to be important, were quantified through analysis of the Nb-peptide interface via computational alanine scan (**Figure 2b**).^36,37^ The consequences of replacing each (non Ala) residue within the peptide epitope with Ala were assessed as a computationally derived ΔΔG value. These computational ΔΔG values were compared between experimental structures and the top ranked Alphafold models for each position within peptide epitopes. If the peptide-Nb interface generated in the AlphaFold model closely resembles the experimental interface there should be a positive and roughly linear correlation when plotting ΔΔG-experimental vs. ΔΔG-model. Analysis of the Nb_Alfa_- and Nb_PepTag_-peptide complexes using this approach yield a positive and linear correlation, whereas other complexes show either a negative correlation, poor concordance between experimental and modeling structures, or both. This analysis shows that the AF2-generated model of the Nb_headlock_ complex places an amino acid residue such that replacement with Ala is predicted to improve binding, suggesting the presence of a steric clash in the AF2 models (**Figure 2b**, red arrows). This position is highlighted in the structural model (**Supporting Figure 1**). Models that impart such energetically unfavorable configurations and interactions are unlikely to be useful for downstream applications.

An analysis of the importance of each residue within the epitope for the Nb-peptide interaction was performed using the AF2 pipeline for Nb_6E_ (**Supporting Figure 2**). Since there are no experimentally determined structures available for Nb_6E_, we compared the properties predicted for AF2 generated models to those previously acquired in experimental studies.^25^ After generating a set of AF2 models of Nb_6E_-6E complexes, we evaluated each of these models using a computational Ala scan. ΔΔG values were calculated for each side chain position within the peptide. As predicted based on visual inspection of the overlaid models, which appeared to show a consistent mode of interaction, variability in ΔΔG values calculated at each position was typically small (**Supporting Figure 2**). Further, the predicted ΔΔG values showed a qualitative parallel with results from analysis of an Ala-scan library of 6E peptides that was previously published (**Supporting Figure 2b**).^25^ Variants of the 6E peptide that showed weak binding in experimental assays mostly possessed Ala mutations at positions that showed high ΔΔG values in the computational Ala scan. One exception is the 6E E5A variant, which showed intermediate binding experimentally and a high ΔΔG value for position 5 in the computational Ala scan. Conversely, Ala mutations that provided 6E variants that showed strong binding experimentally were typically associated with small ΔΔG values. Further evidence for the accuracy and utility of these models of the Nb_6E_-6E complex relates to the favorable confidence metrics calculated by AF2. The iPTM for all five models of the Nb_6E_-6E complex are ≥0.9, which is similar to parameters calculated for the Nb_Alfa_-Alfa complexes that were shown to be highly accurate (**Supporting Table 2**). These findings show that the AF2 models of the Nb_6E_-tag interaction provide a plausible framework for interpreting and predicting the impacts of structural modifications within the 6E tag.

Prior to this work, no experimental structural data was available for the Nb_127_-127 tag complex, even though it has been used as an epitope tag.^32,38^ We experimentally confirmed high affinity binding (K_D_ – 14 nM) between Nb_127_ and the 127 tag peptide using surface plasmon resonance assays (**Supporting Figure 3**). To address this shortcoming and to provide further context for evaluating the performance of AF2 we crystallized the Nb_127_-127 tag complex (**Figure 3a-c)** and using X-ray crystallography solved a 2.1 Å structure. The final model was refined to *R*_work_ = 0.18 and *R*_free_ = 0.22 respectively (**Supporting Table 3**). The asymmetric unit contains a single Nb_127_-127 tag complex in a 1:1 binding mode. The model was clearly defined from Glu1 to Ser115 in Nb_127_, with the 127-tag peptide showing a clear fit within the electron density map (**Figure 3a**). Nb_127_ interacts with the 127-tag via five anti-parallel beta sheets (β4, β5, β6, β9, β10) (**Figure 3b**). This interaction is stabilized by eight strong hydrogen bonds involving residues Glu3’, Phe5’, Trp6’, Lys7’, and Asp10’ of the 127-tag, and Tyr37, Lys96, Lys32, Ser54, and Arg52 of Nb_127_. Water-mediated hydrogen bonds also contribute to the interaction, particularly through a network of four water molecules bridging Gly57 of Nb_127_ with Ser1’ and Phe2’ of the 127-tag (**Figure 3b**). The Nb_127_-127 tag complex is further stabilized by pi-cation interactions between Lys96, Lys32, and Trp6’, as well as pi-pi interactions involving Trp105, Phe2’ and Tyr37, which were observed to reinforce the peptide structure and enhance the engagement between Nb_127_-127 tag complex tag (**Figure 3b**).

We then generated and compared structural models from AF2 for the Nb_127_-127 tag complex to each other and to the experimental data described above. There was a high degree of structural consistency among the AF2 models (**Figure 3c-e**), with only a small divergence in the positioning of the C-terminal residues of the 127 tag (**Figure 3d**, red arrow). Overlay and alignment of the experimental structure with the set of AF2 models also reveals a high degree of similarity (**Figure 3e**). To provide quantitative comparisons of the Nb-tag interfaces seen in AF2 models and in experimental data, we performed computational Ala scans on each of these complexes (**Supporting Table 4, Figure 3f**). There was a high degree of consistency in the predicted energetic contributions (ΔΔG) of 127 tag side chains among the five AF2 models analyzed, resulting in small standard deviations (**Figure 3f**). Very similar trends were observed in assessing the experimental Nb_127_-tag complex structure using a computational Ala scan, with the largest divergences observed for the C-terminal residues (positions 9-10, see **Supporting Table 4**) in the 127-tag peptide. As for the Nb_Alfa_-tag complex described above **(Figure 2a**), we observe a strong positive correlation when plotting ΔΔG values derived from analysis of the experimental structure of the Nb_127_-tag complex vs. ΔΔG values from analysis of an AF2 model (**Figure 3f**).This concordance provides further evidence that AF2 can provide models that are accurate and may be useful predicting the impact of tag modifications on Nb binding.

To test whether the AF2 models or the experimental structural data shown above correlated with experimental binding, we synthesized an Ala-scan library of 127 epitope peptides (see **Supporting Table 5** for sequence information) and assessed their binding to Nb_127_ using a competition-based enzyme-linked immunosorbent assay (ELISA) (**Figure 3g**). In this assay, competitor peptides that bind Nb_127_ more strongly block the binding of Nb to immobilized 127 peptide, resulting in higher levels of inhibition. The findings from these experiments parallel findings from computational analyses of AF2 and experimental structural data (**Figure 3a-c**), which show that important interactions occur between sidechains found at positions 2, 3, 5, and 6 within the 127-tag and its Nb binder (**Figure 3g, Supporting Figure 4**). These parallels provide more support for the utility of AF2 modeling in providing insights into previously structurally uncharacterized Nb-peptide complexes.

To assess whether varying the implementation of AF applied for modeling affected model precision and accuracy, we built and assessed models for each of the Nb-peptide tag pairs using a local version of AF2 (in contrast to the Colabfold running AF2 methodology used elsewhere in the manuscript). Assessments of the local AF2-generated models are shown in **Supporting Table 6**. This new assessment shows many similarities when compared to the modeling performed using ColabFold. Two Nb-peptide tag pair models that closely resemble published experimental data (Nb_Alfa_, Nb_PepTag_) are also associated with high AF confidence scores (model_confidence > 0.85, int_plddt > 90). This correlation further supports the hypothesis that a high AF confidence indicates a higher likelihood of structural accuracy. Models for the other two Nb-peptide tag pairs with published crystal structures were annotated with worse assessment scores (“acceptable” and “incorrect” ratings for Nb_Headlock_ and Nb_Moon_ models, respectively). Application of the local AF2 assessment workflow to the two modeled complexes for which there are no previously reported experimental structures (Nb_6E_ and Nb_127_), also resulted in medium-high confidence scores, consistency with Colabfold generated models, and high similarity with newly generated experimental data (**Supporting Figure 5**). These observations highlight the capability of AF to self-assess the quality of its models to guide further analysis and application of outputs.

We also asked whether AF2 models or experimental structural data would offer useful insights on the rational design of crosslinking analogues of peptide epitopes. Past work^33^ has shown that peptide epitopes linked to electrophiles can rapidly bind and crosslink to a partner Nb, but this approach had not been applied to the Nb-127-tag pair. Peptide compatible electrophiles used in past studies (phenolic esters) are reported to react with Lys and His residues located near to the interface with binding partners.^39^ We sought to use the structural data and models described above to design a peptide-electrophile conjugate that forms a covalent crosslink with its target (Nb_127_). We focused on the incorporation of a crosslinking group at a position within the 127-peptide epitope (Glu3) that was shown (**Figure 3**) to form a hydrogen bond with a Lys residue in Nb (Lys96), which provides the requisite functional group for crosslinking. An analogue of the 127-epitope peptide containing a Cys at position 3 for crosslinker incorporation and a Lys-to-Arg mutation at position 7 (to avoid auto reactivity) was synthesized using conventional solid phase-peptide synthesis (see **Supporting Table 5** for characterization). A crosslinking phenolic ester moiety was attached using Cys-maleimide chemistry as previously described^33^ to provide a peptide-electrophile conjugate (127×3, **Figure 4a**). We then tested this conjugate for its ability to form a covalent crosslink with Nb_127_ as evaluated by mass spectrometry (**Figure 4b**). Mixing Nb_127_ and 127×3 resulted in formation of a new higher molecular weight complex with a mass that corresponded to the formation of the expected crosslinked Nb-peptide product.

**Figure 4.**
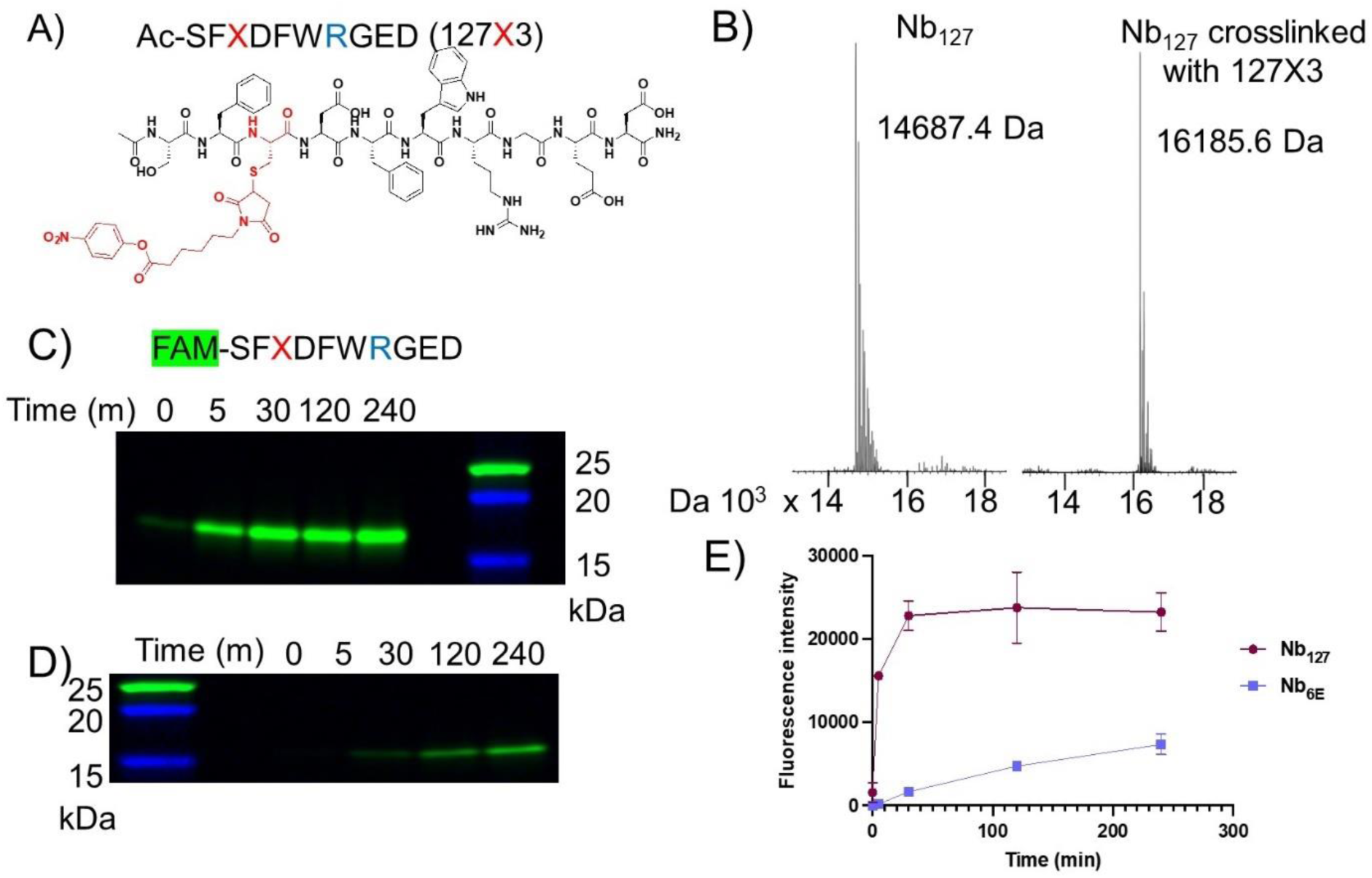
Evaluation of a crosslinking peptide for Nb_127_ guided by modeling and structure. **A)** Structure of an analogue of the 127-peptide tag with a crosslinking group (X) incorporated. **B**) Mass spectrometry analysis of Nb_127_ before (left) and after (right) reaction with Ac-127×3. Crosslinking was performed through mixing Nb_127_ (10 µM) and Ac-127×3 (20 µM) in PBS for 2 h at 25° C followed by purification using a disposable size exclusion column (PD10). **C**) Assessment of the time-dependent formation of a covalently crosslinked complex from Nb_127_ and FAM-127×3 via detection of a fluorescent band on SDS-PAGE. **D**) Assessment of the reaction of FAM-127×3 and Nb_6E_ (negative control) as described in panel c. Fluorescent gel images were obtained as described in **Methods. E**) Quantification of formation of a fluorescent crosslinking product through densitometric analysis of experiments described for panels c-d. Fluorescence intensity was quantified using densitometry as described in **Methods**. The data points are presented as mean ± SD of two independent experiments. An uncropped gel image for data in panels c-d, as well as corresponding Coomassie staining is shown in **Supporting Figure 6**

We then sought to evaluate the kinetics and specificity of crosslinking between Nb_127_ and 127×3. To facilitate easier tracking, we synthesized an analogue of 127×3 containing a carboxyfluorescein fluorophore at its N-terminus (FAM-127×3, **Supporting Table 5**). We evaluated the kinetics of crosslinking between FAM-127×3 and Nb_127_ or Nb_6E_ as a negative control (**Figure 4c-e)**. The mixture of FAM-127×3 and Nb_127_ forms a band corresponding to the crosslinked Nb-peptide product within 5 m, whereas the corresponding reaction with Nb_6E_ is much slower, with a fluorescent band only appearing after 60 m (**Figure 4c-d**). This difference was quantified through densitometric measurement of the fluorescent band intensity (**Figure 4e**). This comparison confirms that the crosslinking performance of 127×3 is accelerated through binding Nb_127_. Despite the rapid initial crosslinking, a qualitative analysis of crosslinking conversion by Coomassie staining shows that crosslinking does not proceed to full conversion (**Supporting Figure 6**). The cause of this stall in crosslinking conversion is unknown and will be investigated in future mechanistic studies.

We also sought to assess whether AF2 modeling could discern between analogues of peptide epitopes that had been shown to bind to Nbs of interest with differing affinity, with analogy to a previously described competition binding approach.^40^ A three-component input consisting of Nb_6E_, 6E-peptide, and 6E-A9 peptide were used to generate a set of models. Past experimental work had shown that Nb_6E_ bound 6E with substantially higher affinity than the 6E-A9 variant.^25^ The models generated by AF2 from the three-component input consistently showed 6E peptide bound at a primary site (1°, **Figure 5a**) that overlays with the site occupied by 6E peptide in models generated from a two-component (Nb_6E_ and 6E) input. In contrast, these three-component models showed 6E-A9 placed in contact with Nb_6E_ at a secondary (2°) binding site that was more variable between models (**Supporting Figure 7**). AF2 generated assessments of confidence in localized protein structures (predicted local distance difference test, pLDDT) also reveal differences between the peptide that binds at the 1° binding site (6E) and the weaker binding peptide (6EA9, **Supporting Figure 7**).

**Figure 5.**
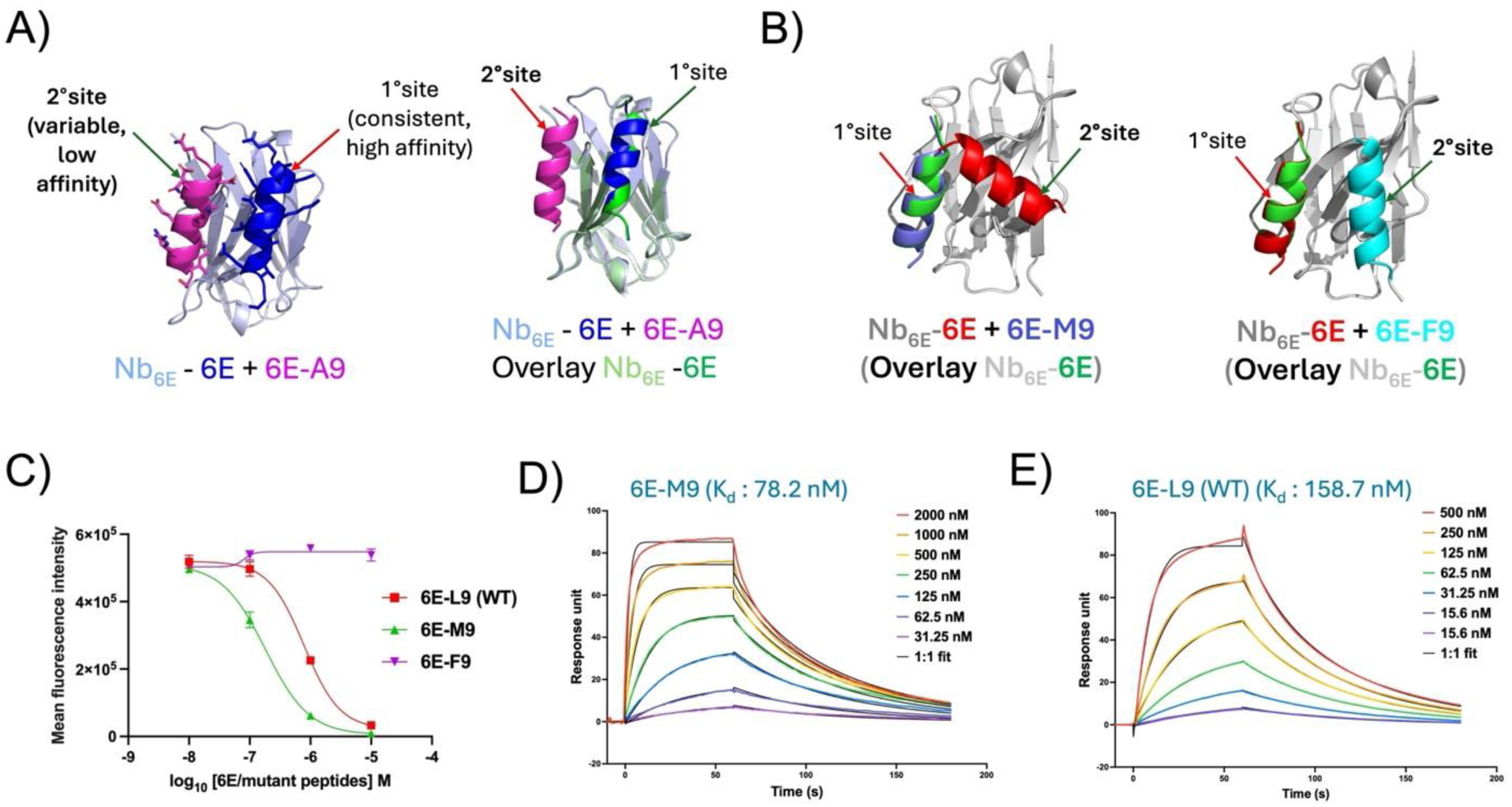
Competition binding using AF2 parallels experimental findings. **A)** (Left) Illustration of primary and secondary binding sites on Nb_6E_ for epitope variants with differing affinities. (Right) Illustration of retention of primary binding site for high affinity epitope (6E) when in competition with a lower affinity variant peptide (6E-A9). **B**) Three component binding models from AF2 predict that one variant peptide (6E-M9, left) outcompetes 6E for binding to the 1° binding site on Nb_6E,_ whereas a different variant peptide (6E-F9, right) does not. **C)** Experimental flow cytometry competition binding assays to assess 6E variant peptide binding. HEK293 cells stably expressing a cell surface protein fused with Nb_6E_ (A2AR-Nb6E-ALFA) were co-incubated with variable concentrations of variant 6E peptides (10 – 0.01 µM) and a fixed concentration (10 nM) of fluorescein-labelled 6E peptide (FAM-6E-C14), followed by washing, and detection with anti-fluorescein antibody conjugated with Alexafluor647 (AF647) (see **Methods**). Binding was quantified as median AF647 fluorescence intensity (see **Supporting Figure 9** for representative histograms). **D-E)** Representative SPR sensorgrams showing association and dissociation curves for interaction of 6E-M9 (2 - 0.03 μM) and 6E-L9 (WT) (0.5 - 0.02 μM) with Nb_6E_-biotin immobilized on a streptavidin (SA) chip (Cytiva, #29104992). Experimental details are described in **Methods**.

We then assessed whether AF2 could be used to predict the relative affinity of novel 6E analogues for Nb_6E_ to prioritize subsequent experimental studies. First, 6E peptide variants containing modifications at a site (position 9, Leu in WT 6E peptide) that were sensitive to an alanine substitution were evaluated using AF2.^25^ An input consisting of Nb_6E_, 6E-L9 (WT), and 6E-M9 showed that the 6E-M9 variant peptide consistently outcompeted 6E-WT for binding to the primary site on Nb_6E_ (**Figure 5b, Supporting Figure 8**). In contrast, 6E-F9 was uniformly outcompeted by 6E-WT. These predictions were then tested experimentally. The binding affinity of peptides for Nb_6E_ was quantified using a competition binding assay (**Figure 5c, Supporting Figure 9**)^25^ and with surface plasmon resonance (SPR, **Figure 5d**). The results from both the assays are in
agreement and confirm AF2 predictions; the ranks of binding affinities in each assay is: 6E-M9 > 6E-WT > 6E-F9. This finding suggests that future studies to enhance the affinity of 6E for Nb_6E_ may benefit from the use of flexible alkyl or heteroalkyl chains at position 9, while aromatic side chains should be avoided.

We also assessed whether the incorporation of a residue with a large aromatic side chain (tryptophan) would be tolerated at other positions within the 6E peptide that were amenable to Ala substitution (**Supporting Figure 2**). Both AF2 and SPR analyses show that these peptides (6E-W4, 6E-W7, 6E-W8) bind Nb_6E_ with affinity that is improved relative to 6E-WT (**Supporting Figure 10**). In the course of these assessments we also compared 6E-WT to an analogue with three glycines appended at the N-terminus (G_3_-6E), used widely in previous studies.^25,33^ Surprisingly, G_3_-6E was superior to 6E-WT both in competition binding assays and as assessed by SPR (**Supporting Figure 11**). A comparison of 6E-WT vs. G_3_-6E binding to Nb_6E_ by AF2 did not match experimental observations, perhaps related to difficulties in modeling and assessing disordered peptide termini (**Supporting Figure 11**).^41^

The effective modeling of selected Nb-peptide epitope interactions led us to probe what role the protein context had on the success of modeling. We thus assessed whether Nb-peptide epitope complexes modelled successfully in isolation would also be modeled appropriately in the context of full-size antigen-Nb complexes. Recently published assessments of using AF to identify Nb epitopes demonstrated accuracy at a modest level (∼50-70%).^42^ Previously published experimental data shows that both Nb_6E_ and Nb_127_ bind the sequences used in epitope tag development when found in the context of intact protein.^19,24,32,38^ We thus generated models of Nb_6E_ with intact antigen (**Figure 6a**, UBC6e-6E tag), Nb_127_ with UBC6e modified to replace the 6E tag with 127 tag (**Figure 6b**, UBC6e-127 tag), Nb_127_ with intact antigen (**Figure 6c**, CXCR2-127 tag) or Nb_6E_ with CXCR2 modified to replace the 127 tag with the 6E tag (**Figure 6d**, CXCR2-6E tag). Models of the UBC6e-Nb_6E_ complex were not consistent with past experimental data^24^, with Nb_6E_ binding at the face of UBC6e opposite the 6E tag, with substantial variability between models (**Figure 6a, Supporting Figure 12**). This observation was surprising, given the good agreement between AF2-generated models of the Nb_6E_-6E tag complex and experimental findings characterizing this interaction (**Supporting Figure 2**). Complications introduced by modeling a complex with full-size UBC6e instead of the isolated tag are likely to be the source of this loss in performance. Notably, the fragment of UBC6e that corresponds to the 6E tag associates with the UBC6e core and buries residues important for binding Nb_6E_ when modeled in this context (**Supporting Figure 13**). Performing an analogous modeling between Nb_127_ and UBC6e-127 tag provided self-consistent models in which the Nb associated with tag, even in an unnatural context (**Figure 6b**). Modeling the interaction of Nb_127_ with CXCR2-127 tag, the antigen used for immunizations to generate Nb_127_,^43^ showed an acceptable concordance with the models of the Nb_127_-127 tag complex in isolation (**Figure 6c**, right), in line with the effective use of the Nb_127_-tag pair in a variety of biological contexts.^32^ Finally, modeling of Nb_6E_ with CXCR2-6E tag also provided a complex that closely overlaid with the Nb_6E_-6E tag complex in isolation (**Figure 6d**). Overall, AF2 generated self-consistent models in three of four test cases, demonstrating its utility for offering Nb-protein complex models to serve as a starting point for hypothesis generation.

**Figure 6.**
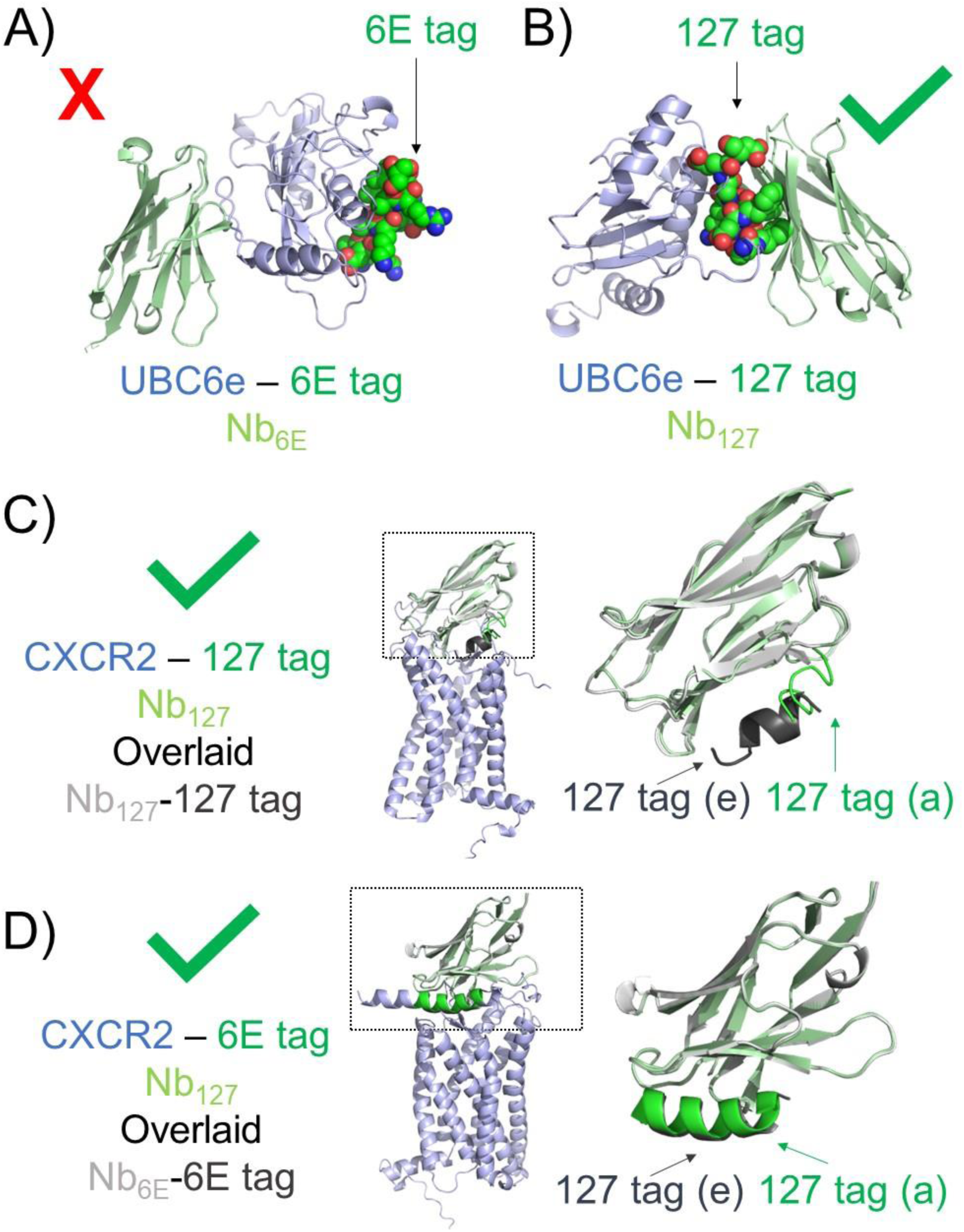
Evaluation of AF2 modeling using full-sized Nb antigens. **A)** AF2 fails to predict experimentally observed binding of Nb_6E_ (light green) to 6E epitope (green spheres, right) when found in the context of UBC6e antigen (light blue). **B)** AF2 successfully predicts binding of Nb_127_ (light green) binding to 127-tag epitope (green spheres) when found in the context of UBC6e antigen (light blue). **C**) AF2 modeling of Nb_127_ (light green) binding to 127-tag epitope (green) when found in the context of full-sized CXCR2 antigen (light blue) shows acceptable overlap with the AF2 model of Nb_127_-127 tag (light gray, dark gray) alone. **D**) AF2 modeling of Nb_6E_ (light green) binding to 6E-tag epitope (green) when found in the context of full-sized CXCR2 antigen (light blue) shows good overlap with the AF2 model of Nb_6E_-6E tag (light gray, dark gray) alone. Insets from dashed boxes (panels c-d) are shown at right. The label “(e)” refers to modeling of the peptide as an epitope and the “(a)” label refers to modeling of the peptide within the full-size antigen. All alignments are performed using Pymol as described in **Methods**.

## Discussion

The release of AF2 and subsequent iterations has furthered scientists’ capability to predict the structure of proteins and protein complexes.^5^ Its adaptation has been empowered by the development of tools, such as the ColabFold platform^34^, that allow users without extensive modeling expertise to generate models that lead to new hypotheses and research directions. Despite widespread adaptation, one of the least successful areas for AF prediction (and predictions of protein-protein interactions more generally) relates to its modelling of antibody-antigen structures. The causes for this challenge are multifaceted; intense and ongoing studies are underway to improve performance in this area.^7–9^ In this work, we perform a focused assessment of the performance of AF2 for generating useful and accurate models of Nbs bound to their targets. Previously published, broad assessments of AF2 performance have evaluated its performance for some Nb-antigen complexes.^8,42^ Here we have delved into this analysis, structurally characterized a novel Nb-epitope interaction, and have tested the utility of AF2 models for creating crosslinking Nb-peptide epitope pairs.

There are several previous studies that have focused on the application of AF for modeling the binding of small peptides to full-sized protein targets^40,44–48^, as well as the binding of antibodies to antigens^8,9,42,49^. In one example, AF2 has been used to model the binding of small peptides to G protein-coupled receptors.^46^ Recent work in the Critical Assessment of Predicted Interactions (CAPRI) initiative has provided insights into the performance of AF2 for modeling antibody-peptide (and antibody-protein antigen) interactions, showing that strategies such as additional AF2 sampling can be helpful, while accurate scoring and model selection is an ongoing challenge.^50–52^ Initially, the success of AF2 in modeling such interactions was surprising, given that most of these small peptides do not adopt a folded structure in solution. However, peptide binding and folding often occur together. As such, these interactions are analogous to interactions between distant sequences within individual proteins, which are effectively modeled by AF2. In this context, it is interesting to note that most of the Nb-tag pairs modelled effectively (Alfa, 127, 6E, and PepTag) are comprised of a peptide epitope that adopts (at least partially) a regular secondary structure (alpha helix). This alone does not guarantee success, as another Nb-bound epitope peptide (Moon) adopts a partially helical conformation upon binding but is modeled incorrectly. We also found that the protein context in which the epitope is situated affects modeling success. Modelling of Nb_6E_ with its 6E peptide epitope alone provides a model consistent with experimentally observed trends. In contrast, modelling Nb_6E_ with 6E epitope found in the context of full-sized antigen (UBC6e) generates models inconsistent with experimental findings. Future efforts to model antibody-antigen (or protein-peptide) interactions may benefit from comparing models produced using full-sized antigens to those generated with isolated epitopes, when possible.

In this work, we find AF2 to be modestly successful in generating accurate and useful models of Nb-peptide epitope complexes. For 4 out of 6 examples, AF2 generates models that either closely mimic experimentally derived structural data or are in good concordance with experimental binding data (**Supporting Tables 1 and 5**). For the other examples, AF2 produces models that are not internally consistent and do not conform with experimental data. In this context, it is worth noting that both Nb-tag pairs that were poorly modeled by AF2 were included in the training set. Memorization of the training set by AF2 is thus no guarantee that models of complexes included in the training will be accurate. Recent studies have shown that even when AF2 models show modest divergences with experimental structural data, they can still be useful for docking campaigns to identify new binders.^53^ It is possible that the AF2 models generated in this work that diverge from experimental structures could still be useful, although further studies will be required to assess this possibility. The determinants of success for generating models of Nb-peptide tag interactions that resemble experimental findings remain unclear. One potentially noteworthy observation relates to a correlation between the length of Nb CDR3 loops, which correspond to the most diverse portion of Nbs sequences, and modelling success. The two Nb-tag pairs modelled least successfully (Nb_Moon_ and Nb_Headlock_) are comprised of Nbs with the longest CDR3 regions (14 and 13 residues, respectively). However, it is not possible to make any firm conclusions on the relationship between CDR3 length and modeling success from this small data set. In summary, these findings demonstrate that AF2 modeling can generate precise and accurate models of Nb-tag interactions, which can be leveraged to design new analogues of epitopes with useful properties such as covalent crosslinking capacity.

## Methods

### Modeling protein-protein complexes using Alphafold

Modeling was performed with AlphaFold2 using the online tool ColabFold (v1.5.5)^34^, as well as the full downloaded AlphaFold2 pipeline. For ColabFold, the Alphafold2_multimer_v3 model was applied for prediction of protein complex structures. Following default ColabFold AlphaFold2 settings, output models were not subjected to relaxation, and no structural templates were used. The MSA mode was mmseqs2_uniref_env. The “unpaired_paired” mode was used in which sequences from the same species and unpaired MSA were paired. A maximum of 3 recycles were allowed along with the “auto” setting for early stop recycle tolerance. The pairing strategy was set to the “greedy” setting. The “max_msa” setting was placed on “auto” and the number of seeds set to “1”. Bipartite and tripartite inputs were submitted with “:” separating each separate polypeptide chain.

The full AlphaFold2 pipeline was downloaded from Github and installed on a local computer cluster and run with the AlphaFold-Multimer v2.2 model. To avoid overlap of templates with modeling targets, a template date cutoff of 2016-04-06 was used for the Nb_PepTag_ and Nb_Headlock_ complexes (corresponding to the Nb_Headlock_ complex, PDB 5IVN, release date), while a template date cutoff of 2018-04-30 was used for the remaining complexes. 25 models were generated per complex, and the top-ranked model based on AlphaFold2 confidence score was assessed. Models were relaxed using the AlphaFold2 Amber relax protocol.

### Generation of input sequences for AlphaFold modeling

Nanobody, epitope tag, and protein sequences were extracted either from the Uniprot database, the Protein Data Bank, or from published sequences. Full sequences for each nanobody and epitope tag used in modeling experiments are found in Supporting Information. Modeling experiments using full length proteins were truncated as listed either in Figure captions or in Supporting Information. Chimeric sequences in which epitope tags are swapped into heterologous full-length proteins are also listed in Supporting Information.

### Structure model visualization using Pymol

Structure models generated by AF2 were visualized using Pymol (version 2.4.1). Alignments of AF2 outputs were performed using the “Align all to this” command. Visualizations were created showing cartoon, surface, or mesh depictions as described in figure captions.

### Model accuracy assessment

Model accuracy metrics were computed based on comparison of modeled complexes with the corresponding X-ray structures using the DockQ program.^54^ Calculated metrics include interface RMSD (I-RMSD), corresponding to backbone RMSD of modeled versus X-ray structure residues at the nanobody-peptide interface, peptide RMSD, corresponding to backbone RMSD of modeled peptide versus X-ray structure peptide after superposition of the nanobody, and CAPRI accuracy level.^35^ CAPRI model accuracy level is based on a combination of I-RMSD, ligand RMSD (peptide RMSD in this context), and fraction of native contacts in the modeled complex, and classifies each model into one of four accuracy levels: Incorrect, Acceptable, Medium, and High. The protein-peptide complex CAPRI criteria were used for assessment of these complexes (“-capri_peptide” flag in DockQ).^35^

### Computational alanine scan to assess contributions of peptide tag side chains to binding

PDB files generated either as AF2 outputs or retrieved from the Protein Data Bank were used as inputs for into the online BUDE Alanine Scan tool.^36,37^ Experimental data were processed using Pymol to remove water molecules prior to input into the online tool, but no other pre-processing was performed. Output ΔΔG values were used to generate graphics and tables shown in figures as described in legends.

### ELISA analysis of Nb_127_ binding by 127-tag peptide analogues

Immobilization of 127-tag peptide was performed by addition and overnight incubation of 127-tag peptide conjugated to green fluorescent protein (using sortagging, see below) on Nunc Maxisorp 96-well plates at a dose of 100 ng/well. Wells were then washed with phosphate buffered saline with 0.05% (v/v) tween-20 (PBST). Wells were then treated with a mixture of Nb_127_-biotin (60 nM, prepared using sortagging) and competitor peptides (concentration 100 μM) for 1 h. Following incubation, all wells were washed 3x with PBST and streptavidin-HRP (Biolegend #405210, 1:2000 dilution) was added for 30 minutes. Washing with PBST (3x) was performed and the plate was treated TMB ELISA solution (Thermo Fisher, #34028). Development was quenched by addition of 1 M sulfuric acid and absorbance was recorded at 480 nm.

### Protein labeling via sortagging

Sortagging reactions were performed using Sortase 5M to label target proteins at their C-terminus as previously described.^19^ Reactions were purified with Ni-NTA affinity chromatography and size exclusion chromatography (PD10 columns, Cytiva # 17085101) to yield site-specifically labeled proteins.

### Protein expression and purification

Nb protein sequences from the literature (see **Supporting Information**) were codon optimized for bacterial expression and cloned into a pET26b expression in frame with pelB and His6 sequences using clone EZ service from GenScript. The production and purification of Nbs has been described previously.^19^ The identity of the purified Nbs was confirmed by mass spectrometry, and the concentration of Nbs was determined by measuring the absorbance at 280 nm.

### Mass spectrometry characterization of purified peptides and proteins

Mass spectrometry data was acquired on a Waters Xevo qTOF LC/MS instrument in positive ion mode. For proteins analyzed by mass spectrometry singly charged ions were not observed, so protein intact mass was calculated from analysis of multiply charged ions using the MaxENT algorithm on MassLynx (Waters) software.

### Peptide synthesis, purification, and modification with a crosslinking group

Alanine scan peptide analogues of 127-tag were ordered via custom synthesis from Genscript and purified before further use (see below). All peptides used for construction of crosslinking conjugates were synthesized in house via solid phase peptide synthesis with Fmoc protection of the amine backbone on a Gyros PurePep Chorus or Liberty Blue Microwave-Assisted Automated Peptide Synthesizer. Peptide synthesis was performed on Rink Amide resin to afford a C-terminal carboxamide. Fmoc-amino acids were dissolved in dimethylformamide (DMF) and added to resin with Oxyma and diisopropylcarbodiimide (DIC). Fmoc groups were deprotected using 10-20% piperidine in DMF. For crosslinking peptides, the N-terminus was acetylated through treatment of washed resin with a mixture of acetic anhydride, diisopropylethylamine, and DMF (1:2:8 v/v) for 10 m.

Synthesized peptides were cleaved from the resin using a cleavage cocktail comprised of trifluoroacetic acid(TFA)/H_2_O/triisopropylsilane(TIS) (92.5:5:2.5% by volume) and rocked at room temperature for 3 hours prior to filtration. Crude peptides were precipitated using diethyl ether. Peptides were purified via preparative-scale HPLC using a Phenomenex Aeris Peptide XB-C18 Prep column (particle size 5 µM, 100 Å pore size) with a linear gradient of solvent A (0.1% TFA in H_2_O) and solvent B (0.1% TFA in acetonitrile). Fractions containing peptides of interest were identified using mass spectrometry analysis. Mass spectrometry characterization of peptides is shown in **Supporting Table 5**.

Fluorescein was appended to the peptide N-terminus on resin, synthesized as described above. To the solid support containing crude, protected peptide, 10 equivalents of 5(6)-Carboxyfluorescein (Acros Organics), 10 equivalents of hexafluorophosphate azabenzotriazole tetramethyl uranium (HATU) and 20 equivalents of diisopropylethylamine were added. The reaction was shaken overnight and washed with DMF. Peptide cleavage and purification was performed as described above.

Peptide electrophile (p-nitrophenol ester) conjugates were synthesized as previously described.^33^ Briefly, purified peptides containing Cys residues were dissolved in DMSO (10 mM) and mixed with 3-5 equivalents of maleimide-phenol ester, which was produced according to a published protocol.^33^ Concentrated phosphate buffer (1 M, pH 7.5) was then added (final concentration 100 mM). The reaction was shaken at room temperature for 1 h and the product was purified by reverse-phase HPLC (C18 column, gradient 20-90% acetonitrile in water with 0.1% trifluoroacetic acid), lyophilized and dissolved in DMSO (1 mM stock) prior to use. Conjugate identity was confirmed via mass spectrometry (**Supporting Table 5**).

### Peptide crosslinking and assessment by SDS-PAGE

Crosslinking reactions were performed using 1 equivalent of nanobody (10 µM) and 2 equivalents of 127×3 crosslinking peptide (20 µM) in PBS at 25 °C for 2 h. The reaction mixture was then purified using fractionation with a size exclusion PD-10 column (Cytiva # 17085101). Fractions containing nanobody were concentrated using a 10 KDa molecular weight cutoff centrifugal filter (Sigma Aldrich, UFC9010). Reaction progress was monitored by SDS-PAGE and mass spectrometry.

The reaction of Nb_127_ and FAM-127×3 was performed as described above, with the following modifications. The reaction was monitored over 4 h. Aliquots were collected at different time points (0, 5, 30, 120 and 240 m) and quenched using a solution of 100 mM DTT in 4x Laemmli sample buffer (BioRad, Cat# 1610747) followed by heating at 95° C for 10 m. Denatured samples were resolved using electrophoresis on a 4-20% acrylamide gradient SDS-PAGE gel (BioRad; Cat# 4561093). The gel was first scanned to detect fluorescence signals, followed by total protein staining using Coomassie stain (Protein Gel Stain, Biotium, Cat# 21003). Fluorescence scans were obtained using the Flamingo (590/110, Blue Epi/ UV filter) application and merged with a Coomassie Blue Far Red Epi application (715/30 Filter) in the ChemiDoc™ MP system (BioRad).

Images were processed for quantification using FiJi software (ImageJ) and by selecting an equal rectangular area for all the fluorescent bands. The mean and standard deviation for the integrated signal for each band were calculated from two independent experiments.

### X-ray crystallography

A fresh preparation of Nb was expressed and purified prior to crystallization. The Nb (5 mg/mL) was mixed with its corresponding peptide epitope tag at molar ratios of 1:2 and 1:3, respectively, at 4°C overnight. The mixture was then centrifuged at high speed to remove debris and any unbound Nb. The crystallization screening was performed using an Art Robbins Gryphon crystallization robot by hanging-drop vapor diffusion method. The screening was performed in a 96-well format (192 conditions) using the Index HT (Hampton Research #HRT-144) crystallization kit. 1 μL Nb-peptide was mixed with 1 μL reservoir solution and then equilibrated against 50 μL reservoir solution and plates were stored at 291 K. Crystal drops were observed daily using a bright-field microscopy. Diffraction-quality crystals of the Nb_127_-127 tag complex were obtained in condition containing 0.1 M Tris pH 8.5, 2.0 M Ammonium sulphate. Diamond-shaped crystals appeared after 3 days equilibration against the crystallization solution and grew to full size of 0.3 × 0.4 × 0.5 mm in 7 days.

### Data collection and structure determination

For data collection, the crystals were briefly soaked in reservoir solution supplemented with 25% (v/v) ethylene glycol and then flash-cooled in liquid nitrogen. X-ray diffraction data were collected at the NIDDK Molecular Structure Facility using a Rigaku Raxis-IV+ and EIGER 4M pixel array detector. Diffraction data were processed using the HKL2000 package.^55^ The crystal structure described in this study was solved by molecular replacement using the model of PDB accession code 6B20 (chain E) as a search model and model building used AutoBuild in Phenix1-1.20.1.^56^ After several cycles of manual adjustments with Coot 1.0^57^, the model was refined with Phenix giving a final R and R_free_ of 0.18 and 0.22, respectively; structural statistics are reported in **Supporting Table 3**. All structure figures were generated using PyMOL 2.3.2 (The PyMOL Molecular Graphics System, Schrodinger). The coordinates and structure factors have been deposited in the Protein Data Bank (PDB) with accession code 9NK9.

### Flow cytometry analysis of peptide binding to HEK293 cells

HEK293 cells stably expressing a cell surface protein fused with Nb_6E_ (A2AR-Nb_6E_-ALFA, described previously^25^) were cultured in DMEM media containing 10% fetal bovine serum (Thermo# A5256801) and 1X pen-strep (Thermo# 15140-122). Binding assays were performed as described previously.^25^ Briefly, cells were suspended in 2% BSA in 1X PBS (w/v) (BSA-PBS) with variable concentrations of unlabeled variant 6E peptides (10 – 0.01 µM) and 10 nM fluorescein-labelled 6E peptide (FAM-6E-C14) for 30 m followed by 2x washing with BSA-PBS, and subsequent staining with anti-fluorescein antibody conjugated with Alexafluor647 (1.5:1000 dilution in BSA-PBS; Jackson ImmunoResearch# 200-602-037). Cells were incubated again for 30 m, pelleted, washed, and analyzed by flow cytometry on a CytoFlex flow cytometer (Beckman Coulter). Cell labelling intensity was monitored in the FL4-APC channel. Live cells were identified and gated based on forward scatter-side scatter profile. A set of 2,000 events corresponding to live cells were captured. Median fluorescence intensity (MFI) for cell counts observed in the APC channel for each sample was calculated from histograms. MFI values obtained from technical replicates of the same sample were averaged and plotted against the corresponding concentration of the variant 6E peptide.

### Surface plasmon resonance analysis (SPR) for Nb-tag binding affinity

SPR biosensor analysis was conducted on a Biacore T200 instrument (Cytiva) using a Series S Sensor Chip SA (streptavidin; Cytiva #29104992)) to determine Nb-tag binding affinity. All binding analyses were done in degassed SPR running buffer (1X PBS at pH 7.4 plus 0.05% P20). Biotinylated Nbs (Nb_6E_ or Nb_127_), produced by sortagging reaction, was captured on SA chip where streptavidin is covalently coupled to dextran matrix. Biotinylated Nb (1 μM) was immobilized on flow cell via biotin-streptavidin interaction to reach a target response value of approximately 2000 response units (RU). A separate flow cell was capped with only SPR running buffer to use as a surface control for nonspecific binding.

Analyte samples (127 tag peptide or mutant-6E peptides) were prepared as serial dilutions in SPR running buffer ranging from 400 - 0.3 nM (or 2 – 0.03 μM). Analyte was flowed over the chip at 50 μL/min with a contact time of 60 seconds, and a dissociation time of 120 sec. A duplicate injection of one concentration and a buffer injection (blank) were used for assessing chip surface performance and double referencing, respectively. The regeneration step (dissociation of bound analyte) included consecutive washes with 10 mM glycine solution (pH 1.5), followed by running buffer for reconditioning. Kinetic constants and K_D_ values (M) were calculated from local fittings using 1:1 kinetics binding model for peptide (analyte) binding to Nb-biotin on the BIAevaluation software (Cytiva).

## Data availability

All data describing the results of this paper are available either within the manuscript, the supporting information, or in the PDB entry described above.

## Conflict of interest

The authors declare that they have no conflicts of interest with the contents of this article.

## Acknowledgements

We acknowledge the intramural mass spectrometry core facility in NIDDK (J. Lloyd) for characterization of peptides and conjugates. This work was supported by the NIH Intramural Research Program (NIDDK, 1ZIADK075157 and 1ZIADK075184 to R.W.C.; ZIADK032103 to C.A.B), and funding from the NIH Director’s Award. This work was also supported by NIH grant GM144083 (to B.G.P.)

## Notes

### Competing Interest Statement

The authors have declared no competing interest.

### Summary of Updates

Graphics in Figure 5 and Supporting Figures 3 + 10 updated to improve clarity

