## Supporting Information for "Evaluation of Alphafold modeling for elucidation of nanobody-peptide epitope interactions"

***Supporting Methods:***

*Input protein sequences*.

$Nb_{6E}$:
QVQLQESGGGLVQPGGSLRLSCAASGFVFENSAMAWYRQAPGKERELIAVIGTTFIKLAESVK GRFTISR DNAKSTVYLQ MNNLKPEDTA VYYCSKSGAYWGQGTQVTVSS

$Nb_{Alfa}$:
EVQLQESGGGLVQPGGSLRLSCTASGVTISALNAMAMGWYRQAPGERRVMVAAVSERGNA MYRESVQGRFTVTRDFTNKMVSLQMDNLKPEDTAVYYCHVLEDRVDSFHDYWGQGTQVTVS S

$Nb_{headlock}$:
QVQLVESGGGLVQPGGSLTLSCTASGFTLDHYDIGWFRQAPGKEREGVSCINNSDDDTYYAD SVKGRFTI FMNNAKDTVYLQMNSLKPEDTAIYYCAEARGCKRGRYEYDFWGQGTQVTV SS

$Nb_{PepTag}$:
QVQLQESGGGLVQPGGSLRLSCAASGNIVSIDAAGWFRQAPGKQREPVATILTGGATNYADS VKGRFTISRDNAKNTVYLQM NSLKPEDTAVYYCYAPMIYY GGRYSDYWGQGTQVTVSS

$Nb_{127}$:
EVQLVESGGGLVQAGESLRLSCAASGSTFDFKVMGWYRQPPGKQREGVAAIRLSGNMHYAE SVKGRFAISKANAKNTVYL QMNSLRPEDTAVYYCKVNIR GQDYWGQGTQVTVSS

$Nb_{Moon}$:
EVQLVESGGGLVQPGGSLRLSCAASGSISSVDVMSWYRQAPGKQRELVAFITDRGRTNYKVS VKGRFTISRDNSKNMVYLQMNSLKPEDTADYLCRAESRTSWSSPSPLDVWGRGTQVTVSS

6E tag: QADQEAKELARQIS

Alfa tag: SRLEEELRRRLTE

BC2 (headlock) tag: PDRVRAVSHWSS

PepTag: AVERYLKDQQLLGIW

127 Tag: SFEDFWKGED

Moon tag: KNEQELLELDKWASL

UBC6E(1-184), 6E tag underlined:
METRYNLKSPAVKRLMKEAAELKDPTDHYHAQPLEDNLFEWHFTVRGPPDSDFDGGVYHGRI VLPPEYPM
KPPSIILLTANGRFEVGKKICLSISGHHPETWQPSWSIRTALLAIIGFMPTKGEGAIGSLDYTPEE RRAL
AKKSQDFCCEGCGSAMKDVLLPLKSGSDSSQADQEAKELARQIS

UBC6E(1-184)-127 tag, tag underlined:

METRYNLKSPAVKRLMKEAAELKDPTDHYHAQPLEDNLFEWHFTVRGPPDSDFDGGVYHGRI
VLPPEYPM
KPPSIILLTANGRFEVGKKICLSISGHHPETWQPSWSIRTALLAIIGFMPTKGEGAIGSLDYTPEE
RRAL
AKKSQDFCCEGCGSAMKDVLLPLKSGSDSSSFEDFWKGED

CXCR2(1-360), 127 tag underlined:
MEDFNMESDSFEDFWKGEDLSNYSYSSTLPPFLLDAAPCEPESLEINKYFVVIIYALVFLLSLLG
NSLVM
LVILYSRVGRSVTDVYLLNLALADLLFALTLPIWAASKVNGWIFGTFLCKVVSLLKEVNFYSGILLL
ACI
SVDRYLAIVHATRTLTQKRYLVKFICLSIWGLSLLLALPVLLFRRTVYSSNVSPACYEDMGNNTA
NWRML
LRILPQSFGFIVPLLIMLFCYGFTLRTLFKAHMGQKHRAMRVIFAVVLIFLLCWLPYNLVLLADTL
MRTQ
VIQETCERRNHIDRALDATEILGILHSCLNPLIYAFIGQKFRHGLLKILAIHGLISKDSLPKDSRPSF
VG
SSSGHTSTTL

CXCR2(1-360)-6E tag, 6E tag underlined:
MEDFNMESDQADQEAKELARQISLSNYSYSSTLPPFLLDAAPCEPESLEINKYFVVIIYALVFLLS
LLGNSLVM
LVILYSRVGRSVTDVYLLNLALADLLFALTLPIWAASKVNGWIFGTFLCKVVSLLKEVNFYSGILLL
ACI
SVDRYLAIVHATRTLTQKRYLVKFICLSIWGLSLLLALPVLLFRRTVYSSNVSPACYEDMGNNTA
NWRML
LRILPQSFGFIVPLLIMLFCYGFTLRTLFKAHMGQKHRAMRVIFAVVLIFLLCWLPYNLVLLADTL
MRTQ
VIQETCERRNHIDRALDATEILGILHSCLNPLIYAFIGQKFRHGLLKILAIHGLISKDSLPKDSRPSF
VG
SSSGHTSTTL

### Supporting Tables.

### Supporting Table 1. Summary of nanobodies and epitopes used in modeling studies.

| Nanobody name | Structure (PDB code) | Release date | Pubmed ID(s) | Tag name | Tag sequence | Modelling success summary |
|---|---|---|---|---|---|---|
| VHH05 /VHH6E/ Nb6E | None | - | 35930411, 31518855 | 6E tag | QADQEAKELARQIS (ubc6e) | Predicts important residues |
| Alfa | 6I2G | 9/25/2019 | 31562305 | Alfa | SRLEEELRRRLTE | Very good match |
| Headlock | 5IVN | 4/6/2016 | 26791954, 29500346 | BC2 | PDRVRAVSHWSS (beta catenin) | Correct general site, some divergence in details |
| Peptag Nb, 2E7 | 5HM1 (no paper) | 1/25/2017 | 32868807 | Peptag | AVERYLKDQQLLGIW (from HIV GP41) | Only top model is highly accurate, others diverge |
| 127d01 | None (reported here) | - | 35076390 | CXCR2 ECD | SFEDFWKGED (human CXCR2) | Good match except for peptide C-terminus |
| Moon, 2H10 | 4B50 (no ligand), 7AEJ (full prot) | 3/27/2013, 5/26/2021 | 31178119 | Moon | KNEQELLELDKWASL (from HIV GP41) | Poor prediction |

**Supporting Table 2**. Computational confidence parameters for AF2 models and comparison to experimental structures.

| | | | | | Rank 1 comparison with experimental[1] | | |
|---|---|---|---|---|---|---|---|
| **Complex** | **Rank (ColabFold)** | **pLDDT** | **pTM** | **ipTM** | **I-RMSD, Å** | **Peptide RMSD, Å** | **Model Accuracy** |
| NbAlfa | 1 | 97.2 | 0.917 | 0.913 | 0.33 | 0.46 | High |
| | 2 | 97.2 | 0.917 | 0.912 | | | |
| | 3 | 97 | 0.915 | 0.911 | | | |
| | 4 | 97.2 | 0.917 | 0.909 | | | |
| | 5 | 96.7 | 0.914 | 0.91 | | | |
| NbHeadlock | 1 | 81.1 | 0.814 | 0.409 | 2.28 | 3.62 | Acceptable |
| | 2 | 80.7 | 0.81 | 0.361 | | | |
| | 3 | 80.2 | 0.81 | 0.358 | | | |
| | 4 | 80.3 | 0.806 | 0.269 | | | |
| | 5 | 80 | 0.809 | 0.254 | | | |
| NbPepTag | 1 | 90 | 0.872 | 0.841 | 1.82 | 1.5 | Medium |
| | 2 | 85 | 0.837 | 0.653 | | | |
| | 3 | 86.9 | 0.846 | 0.624 | | | |
| | 4 | 84.7 | 0.826 | 0.501 | | | |
| | 5 | 81.5 | 0.809 | 0.355 | | | |
| NbMoon | 1 | 83.9 | 0.828 | 0.458 | 2.68 | 4.45 | Incorrect |
| | 2 | 86.1 | 0.833 | 0.398 | | | |
| | 3 | 82.4 | 0.811 | 0.381 | | | |
| | 4 | 83.6 | 0.819 | 0.341 | | | |
| | 5 | 82.1 | 0.81 | 0.334 | | | |
| Nb6E | 1 | 97.4 | 0.918 | 0.914 | No experimental structure available | | |
| | 2 | 97.3 | 0.92 | 0.913 | | | |
| | 3 | 97.3 | 0.919 | 0.91 | | | |
| | 4 | 97 | 0.917 | 0.909 | | | |
| | 5 | 96.7 | 0.915 | 0.904 | | | |
| Nb127 | 1 | 92.2 | 0.89 | 0.849 | 0.47 | 1.17 | Medium |
| | 2 | 92.6 | 0.891 | 0.841 | | | |
| | 3 | 92.5 | 0.894 | 0.837 | | | |
| | 4 | 92.8 | 0.896 | 0.836 | | | |
| | 5 | 92.5 | 0.894 | 0.821 | | | |

[1]Model accuracy assessments for interface residue RMSD (I-RMSD), peptide RMSD, and CAPRI model accuracy level were computed based on comparison of model with corresponding X-ray structure using the DockQ program.[1]

**Supporting Table 3**. Complete data and statistics (diffraction data and refinement statistics) for X-ray analysis of $Nb_{127}$-127 tag complex crystal.

| Data collection | $Nb_{127}$ |
|---|---|
| ligand | 127D01 |
| Space group | P 31 2 1 |
| Wavelength (Å) | 1.54 |
| Resolution (Å) | 50-2.10 |
| Highest resolution shell (Å) | 2.14-2.10 |
| Unit-cell parameters (Å) | a=46.876<br>b=46.876<br>c=91.184 |
| α=β=**γ** (°) | 90, 90, 120 |
| Measurements | |
| Unique reflections | 7180(369) |
| Redundancy | 8.9(6.9) |
| Completeness (%) | 99.3(98.4)[a] |
| *Rmerg*[b] | 0.045(0.104) |
| <I/ $\sigma(I)$> | 51.6(14.1) |
| **Refinement** | |
| No. of reflection (test) | 7149(712) |
| Total atom (non-H) | 1055 |
| No. atoms | |
| Protein | 900 |
| ligand | 89 |
| Water | 76 |
| *R* | 0.18 |
| $R_{free}$ | 0.22 |
| RMSD | |
| Bond lengths(Å) | 0.003 |
| bond angles (°) | 0.60 |
| Average B factor (Å$^2$) | |
| Subunit A | 25.8 |
| Subunit B | 23.9 |
| water | 31.6 |
| Ramachandran plot | |
| Favored (%) | 100 |
| Allowed (%) | 0.00 |
| Outliers (%) | 0.00 |

[a]Figures in brackets apply to the highest-resolution shell.

[b]$R_{merg} = \Sigma_h \Sigma_i | I(h,i) - \langle I(h) \rangle | / \Sigma_h \Sigma_i I(h,i)$, where $I(h,i)$ is the intensity of the *i*th observation of reflection *h*, and $\langle I(h) \rangle$ is the average intensity of redundant measurements of reflection *h*.

**Supporting Table 4: Assessment of side chain contributions to nanobody-tag interactions for $Nb_{127}$.**

| | | Calc. ΔΔG (kJ/mol ± SD) | | | |
|---|---|---|---|---|---|
| **Residue** | **Amino Acid** | **Alphafold structures** | | | **Experimental (xtal strucuture)** |
| 2 | PHE | 14.1 | ± | 1.2 | 16.4 |
| 3 | GLU | 5.5 | ± | 0.2 | 5.7 |
| 4 | ASP | 0.0 | ± | 0.0 | 0 |
| 5 | PHE | 13.7 | ± | 0.1 | 13 |
| 6 | TRP | 27.2 | ± | 0.4 | 25.4 |
| 7 | LYS | 2.0 | ± | 0.1 | 1.7 |
| 9 | GLU | 0.6 | ± | 0.7 | 3.2 |
| 10 | ASP | 5.2 | ± | 4.2 | 9.3 |

**Supporting Table 5. Sequence and mass spec characterization of peptides used in study.**

| Peptide | Sequence | Calculated (M+H) | Observed (M+H) |
|---|---|---|---|
| 127A1 | H-AlaPheGluAspPheTrpLysGlyGluAsp-NH2 | 1242.54 | 1242.5 |
| 127A2 | H-SerAlaGluAspPheTrpLysGlyGluAsp-NH2 | 1182.5 | 1182.5 |
| 127A3 | H-SerPheAlaAspPheTrpLysGlyGluAsp-NH2 | 1200.5 | 1200.5 |
| 127A4 | H-SerPheGluAlaPheTrpLysGlyGluAsp-NH2 | 1214.5 | 1214.6 |
| 127A5 | H-SerPheGluAspAlaTrpLysGlyGluAsp-NH2 | 1182.5 | 1182.5 |
| 127A6 | H-SerPheGluAspPheAlaLysGlyGluAsp-NH2 | 1143.5 | 1143.5 |
| 127A7 | H-SerPheGluAspPheTrpAlaGlyGluAsp-NH2 | 1201.5 | 1201.5 |
| 127A8 | H-SerPheGluAspPheTrpLysGluGluAsp-NH2 | 1330.6 | 1330.6 |
| 127A9 | H-SerPheGluAspPheTrpLysGlyAlaAsp-NH2 | 1200.5 | 1200.5 |
| 127A10 | H-SerPheGluAspPheTrpLysGlyGluAla-NH2 | 1214.5 | 1214.6 |
| 127C3 | Ac-SerPheCysAspPheTrpArgGlyGluAsp-NH2 | 1301.5 | 1302.5 |
| 127X3 | Ac-SerPheCys(xlink)AspPheTrpArgGlyGluAsp-NH2 | 1633.61 | 1634.6 |
| FAM-127C3 | FAM-SerPheCysAspPheTrpArgGlyGluAsp-NH2 | 1617.6 | 1619.6 |
| FAM-127X3 | FAM-SerPheCys(xlink)AspPheTrpArgGlyGluAsp-NH2 | 1949.7 | 976.3 (z=2) |
| 6E-WT | H-GlnAlaAspGlnGluAlaLysGluLeuAlaArgGlnIleSer-NH2 | 794.0 (z=2) | 793.9 (z=2) |
| 6E-M9 | H-GlnAlaAspGlnGluAlaLysGluMetAlaArgGlnIleSer-NH2 | 887.9 (z=2) | 887.9 (z=2) |
| 6E-F9 | H-GlnAlaAspGlnGluAlaLysGluPheAlaArgGlnIleSer-NH2 | 810.9 (z=2) | 810.9 (z=2) |
| 6E-W4 | H-GlnAlaAspTrpGluAlaLysGluLeuAlaArgGlnIleSer-NH2 | 908.4 (z=2) | 908.5 (z=2) |
| 6E-W7 | H-GlnAlaAspGlnGluAlaTrpGluLeuAlaArgGlnIleSer-NH2 | 908.0 (z=2) | 908.0 (z=2) |
| 6E-W8 | H-GlnAlaAspGlnGluAlaLysTrpLeuAlaArgGlnIleSer-NH2 | 907.5 (z=2) | 907.5 (z=2) |
| G3-6E | H-GlyGlyGlyGlnAlaAspGlnGluAlaLysGluLeuAlaArgGlnIleSer-NH2 | 879.0 (z=2) | 879.0 (z=2) |

**Supporting Table 6: Scores and accuracy assessments of Nb-tag complexes using AlphaFold2 run locally.**

| Nanobody name | pTM | ipTM | Model confidence | I-pLDDT | I-RMSD, Å[1] | Peptide RMSD, Å[1] | Model Accuracy[1] |
|---|---|---|---|---|---|---|---|
| VHH05/VHH6E/Nb6E[2] | 0.92 | 0.91 | 0.91 | 97.63 | - | - | - |
| Alfa | 0.92 | 0.92 | 0.92 | 97.91 | 0.4 | 0.61 | High |
| Headlock | 0.85 | 0.49 | 0.56 | 75.46 | 1.68 | 2.17 | Acceptable |
| Peptag Nb, 2E7 | 0.89 | 0.91 | 0.9 | 91.29 | 2.05 | 1.66 | Medium |
| 127d01 | 0.9 | 0.89 | 0.89 | 97.22 | 2.27 | 3.91 | Acceptable |
| Moon, 2H10 | 0.84 | 0.39 | 0.48 | 72.44 | 8.79 | 14.75 | Incorrect |

[1]Model accuracy assessments for interface residue RMSD (I-RMSD), peptide RMSD, and CAPRI model accuracy level were computed based on comparison of model with corresponding X-ray structure using the DockQ program (1).

[2]No X-ray structure is available for this complex, so accuracy assessments were not possible to calculate (noted as “-“).

**Supporting Figures.**

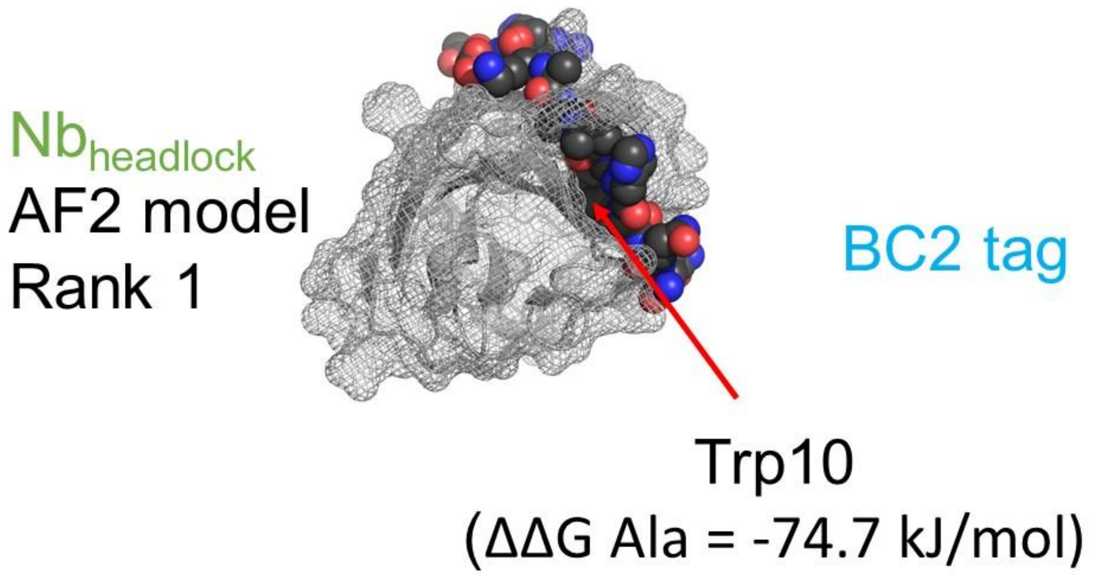


**Supporting Figure 1. Illustration of steric clashes generated in AF2 models.** Top ranking AF2 models for the $Nb_{headlock}$ and $Nb_{PepTag}$ complexes are shown with nanobody depicted in mesh and the binding peptide tag shown as spheres. Residues showing a negative ΔΔG Ala value in computational Ala scan and their steric clashes with the Nb surface are highlighted with arrows.

A) Nb-6E

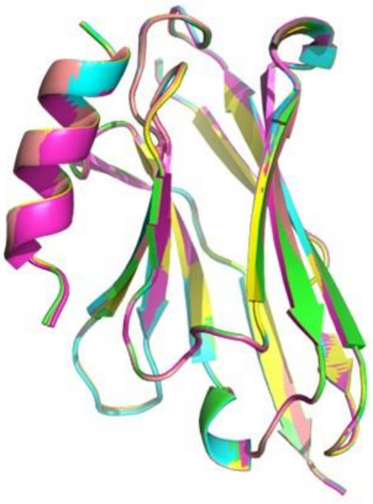

B) Computational analysis Alphafold

| Residue | Amino Acid | ΔΔG (kJ/mol) | | SD |
|---|---|---|---|---|
| 1 | GLN | 2.5 | ± | 0.2 |
| 3 | ASP | 11.9 | ± | 0.3 |
| 4 | GLN | 0.0 | ± | 0.0 |
| 5 | GLU | 12.0 | ± | 0.7 |
| 7 | LYS | 1.6 | ± | 0.0 |
| 8 | GLU | 0.0 | ± | 0.0 |
| 9 | LEU | 8.9 | ± | 0.7 |
| 11 | ARG | 0.1 | ± | 0.0 |
| 12 | GLN | 2.8 | ± | 0.3 |
| 13 | ILE | 9.4 | ± | 0.1 |

Experimental: Large, medium, or small role in binding (Ref Cabalteja *et al.* ACS Chemical Biology 2022)

**Supporting Figure 2. Comparison of AF2 predicted structures to experimental observations.** A) Overlay of five AF2 models generated for the $Nb_{6E}$-6E tag complex. B) Computational Ala scan analysis peptide tag residues contributing to binding of AF2 models. Each of the five AF2 models was analyzed via computational Ala scan. Values shown represent mean ± SD upon analysis of all models.

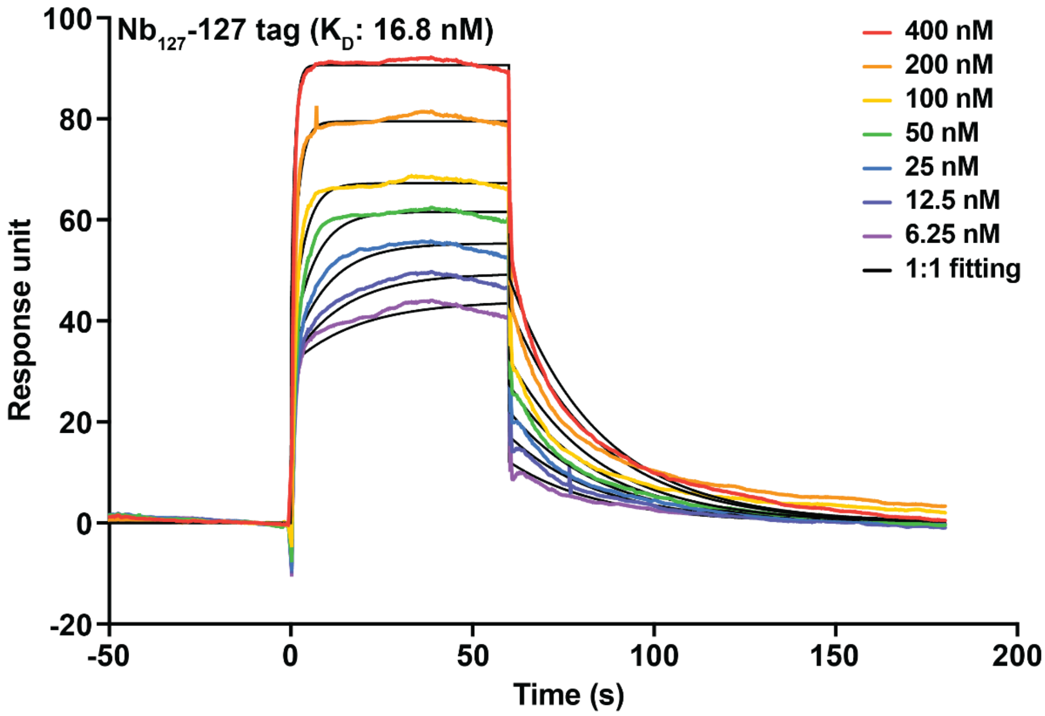


**Supporting Figure 3. Characterization of $Nb_{127}$-127 tag affinity using surface plasmon resonance.** Representative SPR sensorgram showing association and dissociation curves for interaction of 127 tag (400 – 6.25 nM) with $Nb_{127}$-biotin immobilized on a streptavidin (SA) chip (Cytiva, #29104992). Experimental details are described in **Methods**.

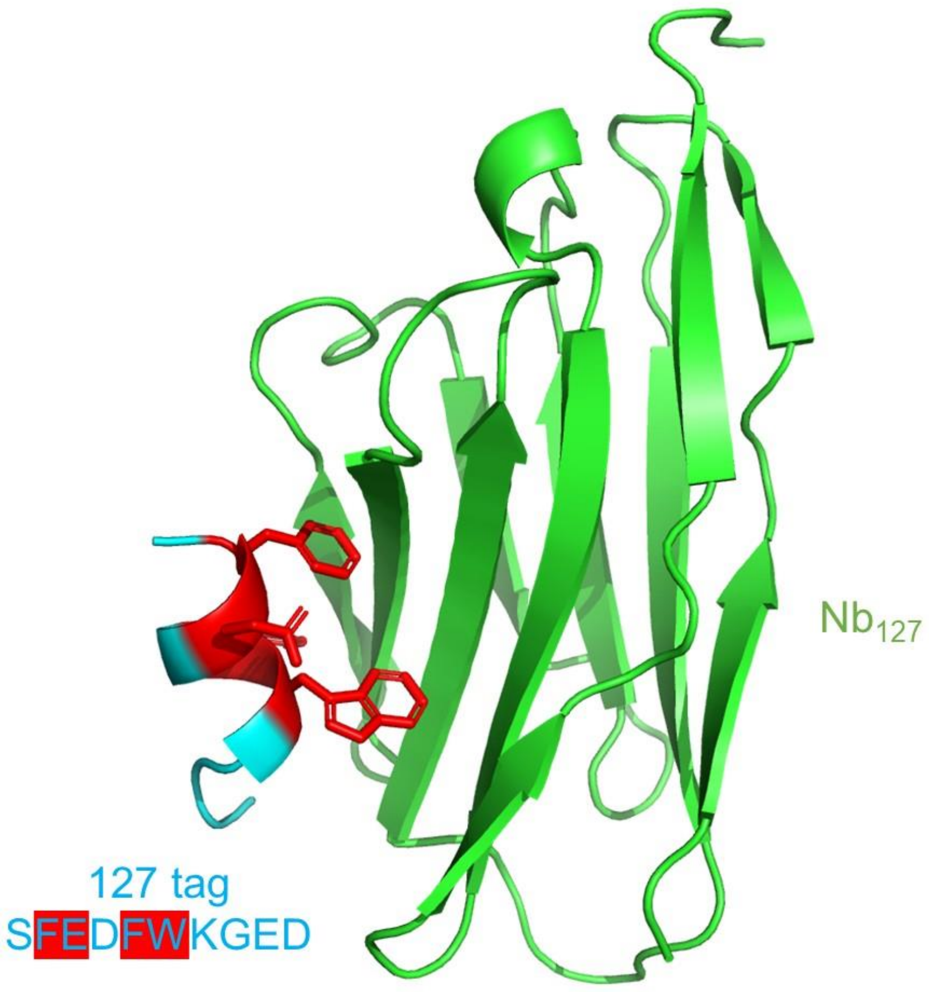


**Supporting Figure 4. Illustration of residues important for binding withing the Nb127-tag experimental structure.** Within the 127-tag peptide (cyan) residues shown to be important for binding in an ELISA are highlighted in red.

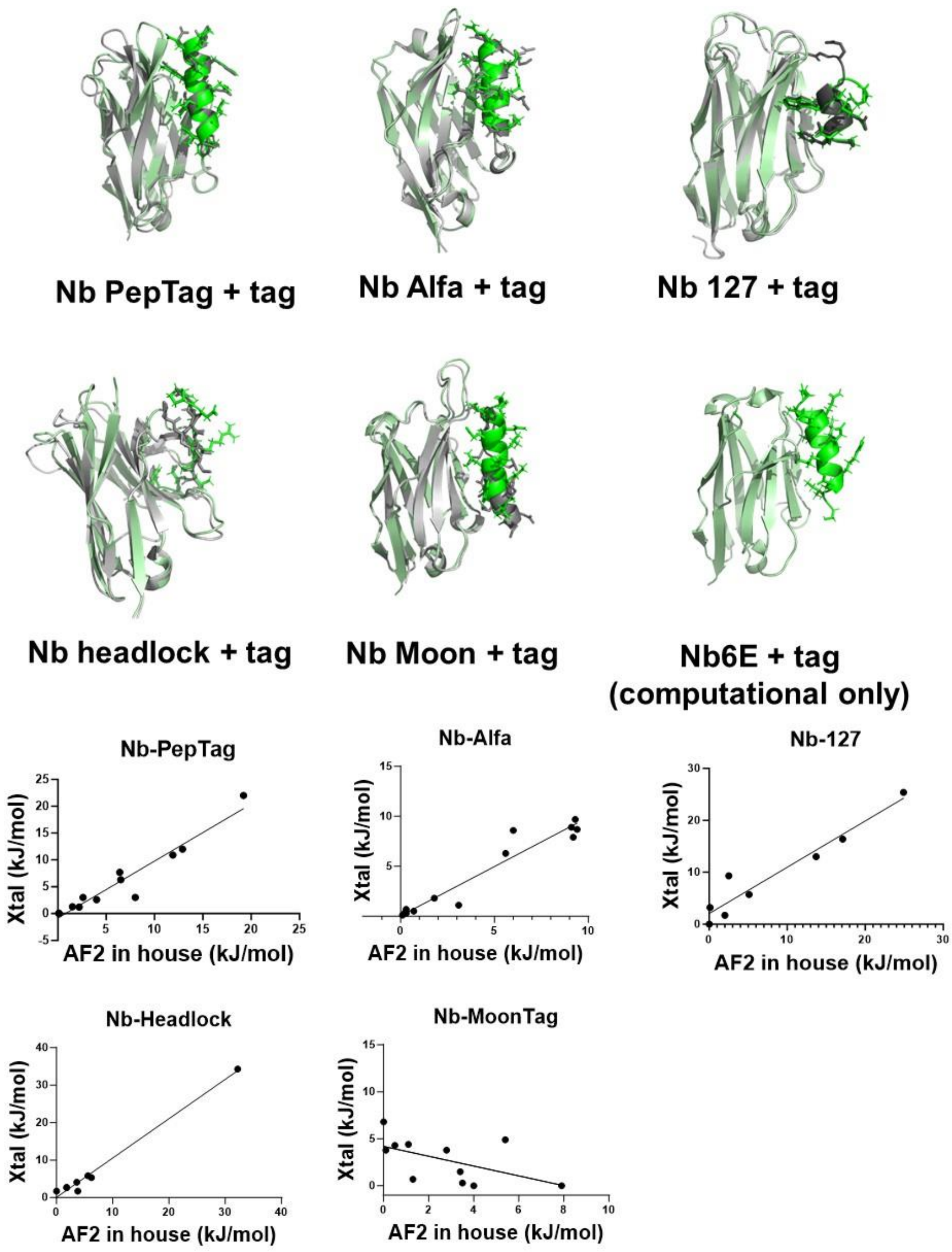


**Supporting Figure 5. Overlay of experimental structures and top ranked models from application of *in house* AF2 implementation.** (Top) Experimental models (Nb in light gray, tag in dark gray) were aligned with top ranked AF2 models (Nb in light green, tag in green) in Pymol. Tag peptides are shown with side chains illustrated with sticks. (Bottom) Quantitative assessment of peptide epitope side chain energetic contributions to binding. The energetic contribution of each side chain within experimental or the top ranked AF2 generated complex was quantified using BUDE Ala Scan. Each side chain was plotted as a single point in a scatter plot.

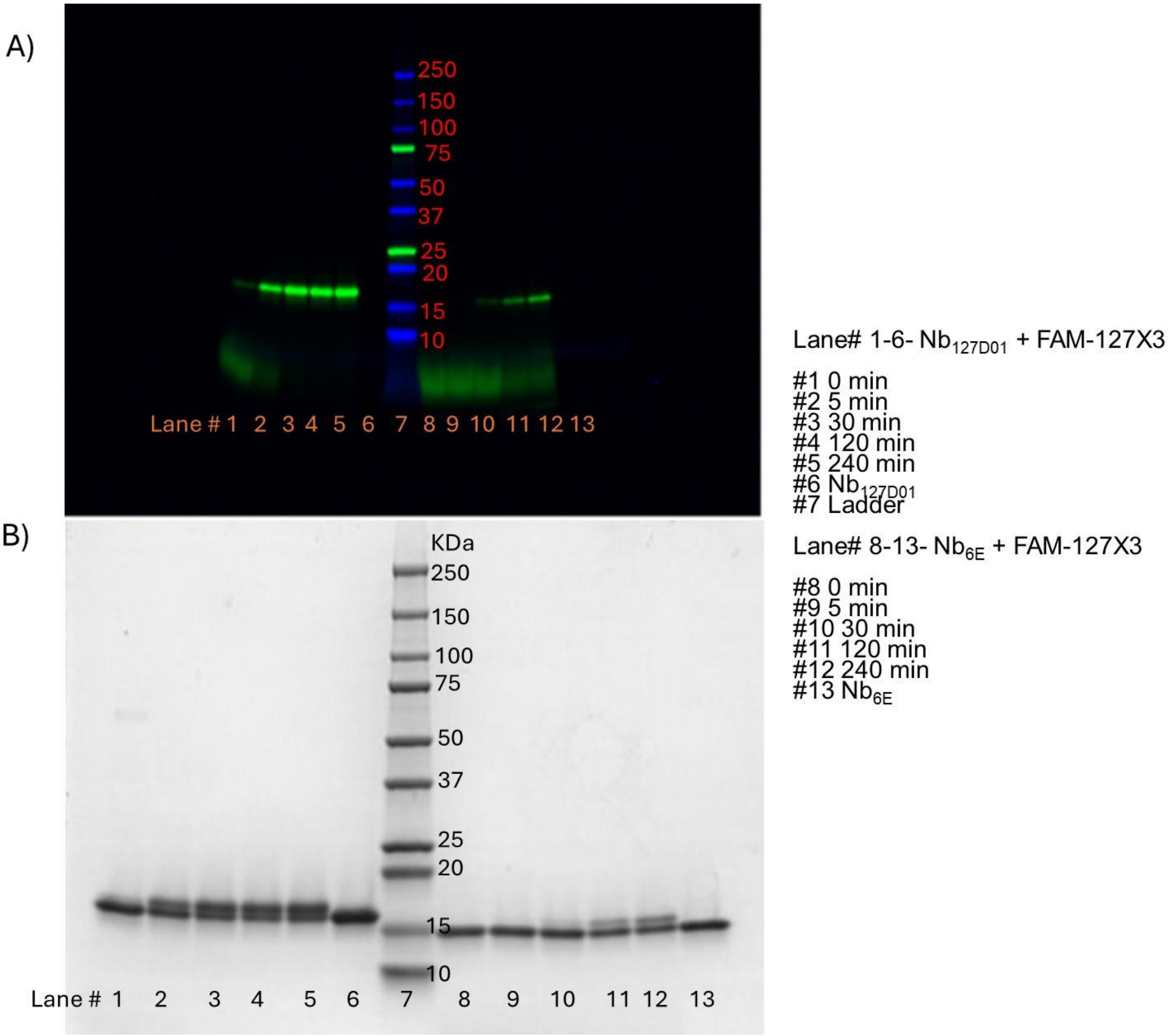


**Supporting Figure 6: Comparison of the labeling of $Nb_{127}$ and $Nb_{6E}$ with FAM-127X3.** Labeling reactions were performed with a mixture of indicated nanobodies (10 µM) and FAM-127X3 peptide (20 µM) in PBS at 25° C. The aliquots from the reaction mixture were collected and quenched at indicated time points using 4x SDS-PAGE sample buffer (Biorad # 1610747) containing 100 mM DTT. Reactions were then resolved using electrophoresis on a 4-20% acrylamide gradient gel (BioRad, #4561093). The gel was first scanned to detect the fluorescence (488 nm) and then stained with Coomassie blue for determination of total protein content. Representative gel images of a **A**) fluorescence scan (488 nm) and **B**) Coomassie staining for indicated reactions with conditions described above.

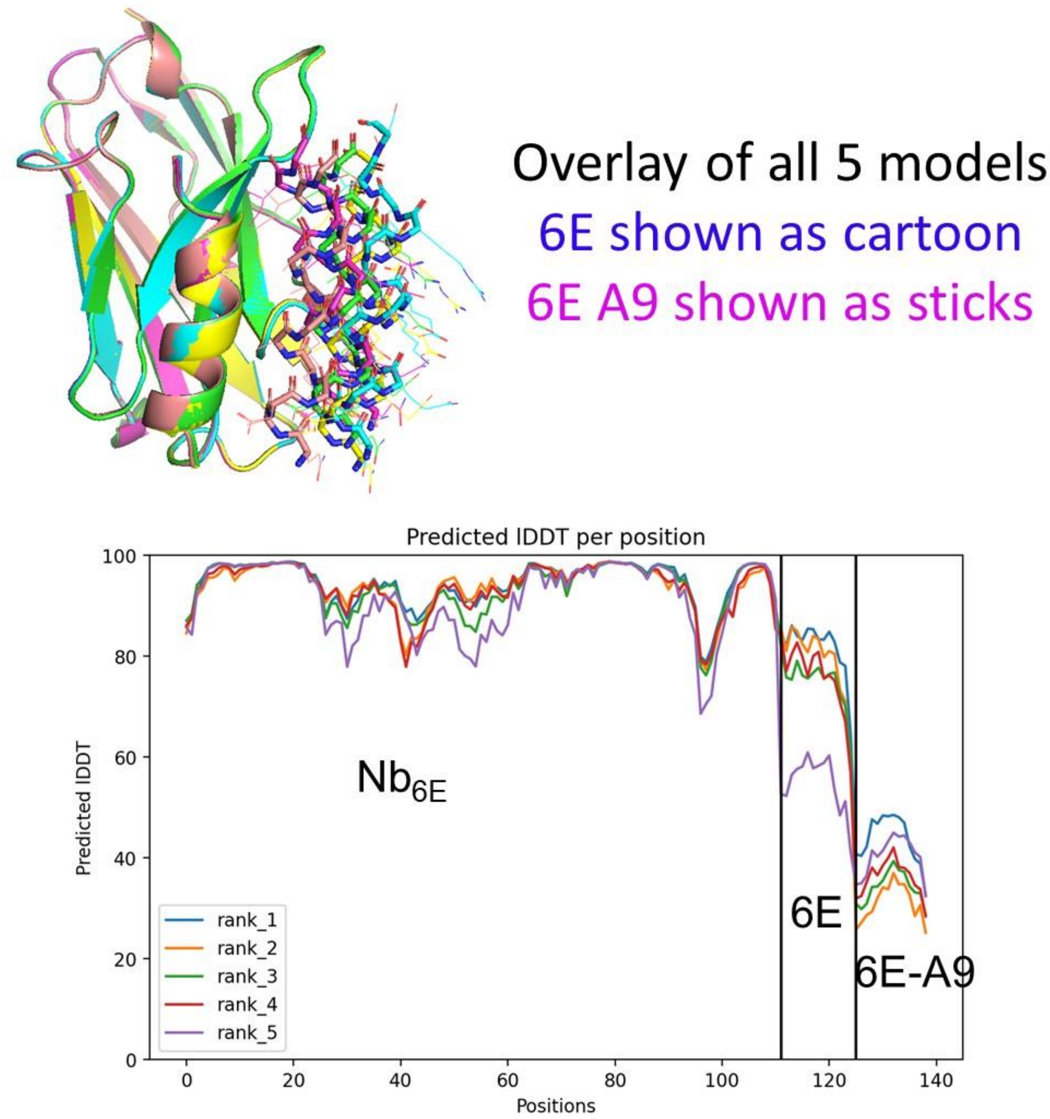


**Supporting Figure 7. Comparison of peptide localization among AF2 models within peptide competition-based (6E vs 6E-A9).** (Upper) An overlay of all 5 AF2 models (each its own color) generated for the tripartite complex consisting of $Nb_{6E}$, 6E, and 6E-A9 is shown. $Nb_{6E}$ and 6E show a high degree of overlap among all models and are shown as cartoons. 6E-A9 shows weaker overlap and is shown as sticks. (Lower) pIDDT plot generated by Colabfold analyzing confidence in local structures across the predicted complex. Note the higher confidence levels shown for 6E relative to 6E-A9.

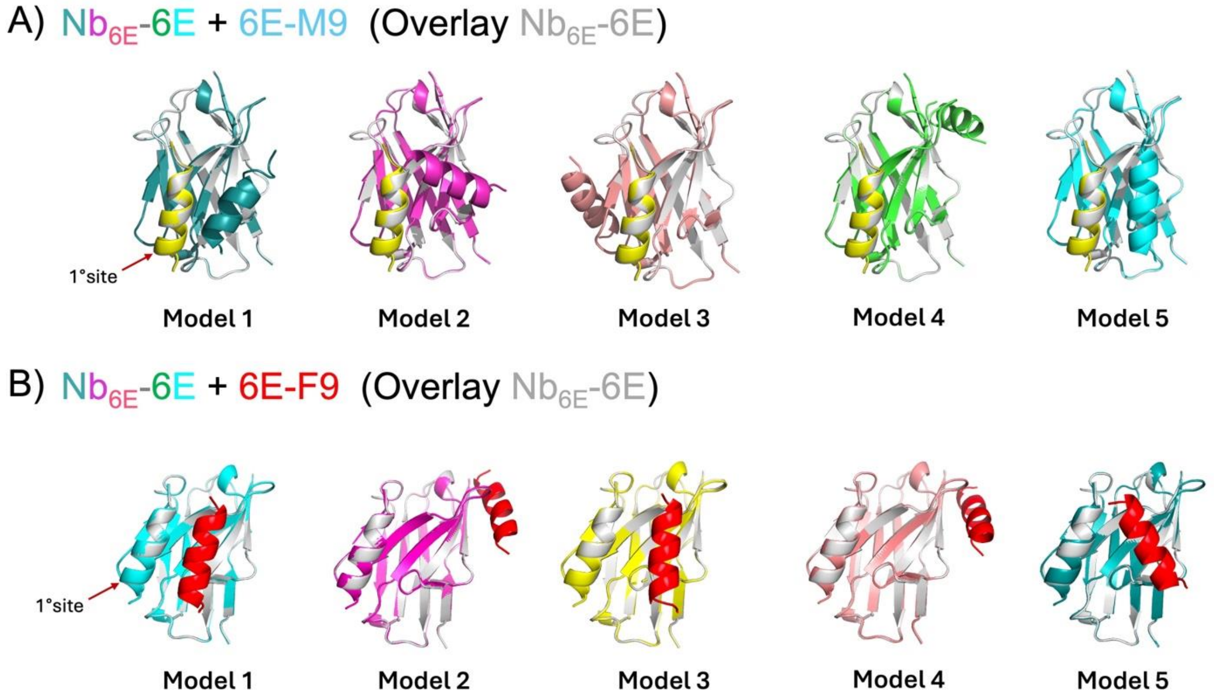


**Supporting Figure 8. Evaluation of binding for 6E mutants via competition-based complex formation using AF2 models.** Individual AF2 generated models were aligned and overlayed with a control model where 6E peptide occupies the primary (1°) binding site on the $Nb_{6E}$ surface (in grey). **A)** 6E-M9 (yellow) outcompetes 6E (multicolor) for the 1° binding site on $Nb_{6E}$ in all models. **B)** 6E-F9 (red) does not outcompete 6E (multicolor) for the 1° binding site on Nb6E.

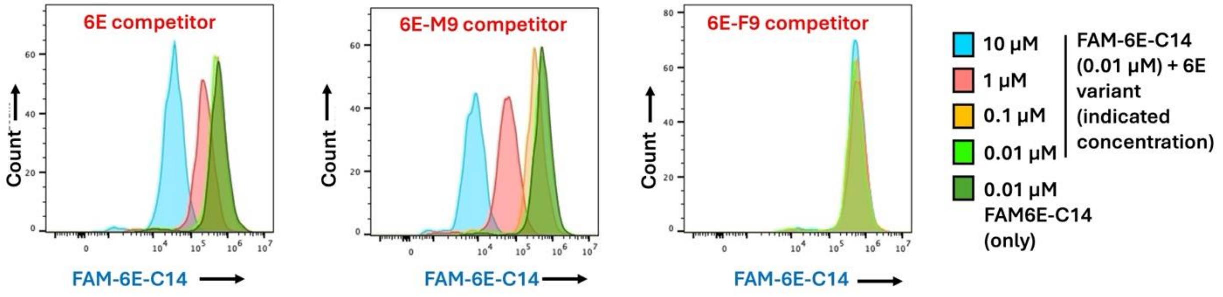


**Supporting Figure 9. Competition binding assays to measure 6E peptide binding.** Representative histograms from flow cytometry assays to measure 6E variant peptide binding to HEK293 cells stably expressing a cell surface protein fused with $Nb_{6E}$ (A2AR-$Nb_{6E}$-ALFA). Cells were co-incubated with variable concentrations of variant 6E peptides (10 – 0.01 $\mu$M) and a fixed concentration (10 nM) of fluorescein-labelled 6E peptide (FAM-6E-C14), followed by washing, and detection with anti-fluorescein antibody conjugated with AF647 (see **Methods**). Histograms depict the intensity of AF647 signaling on live cells from a single representative experiment. These data were used to construct the dose-response shown in **Figure 5c** in the main text.

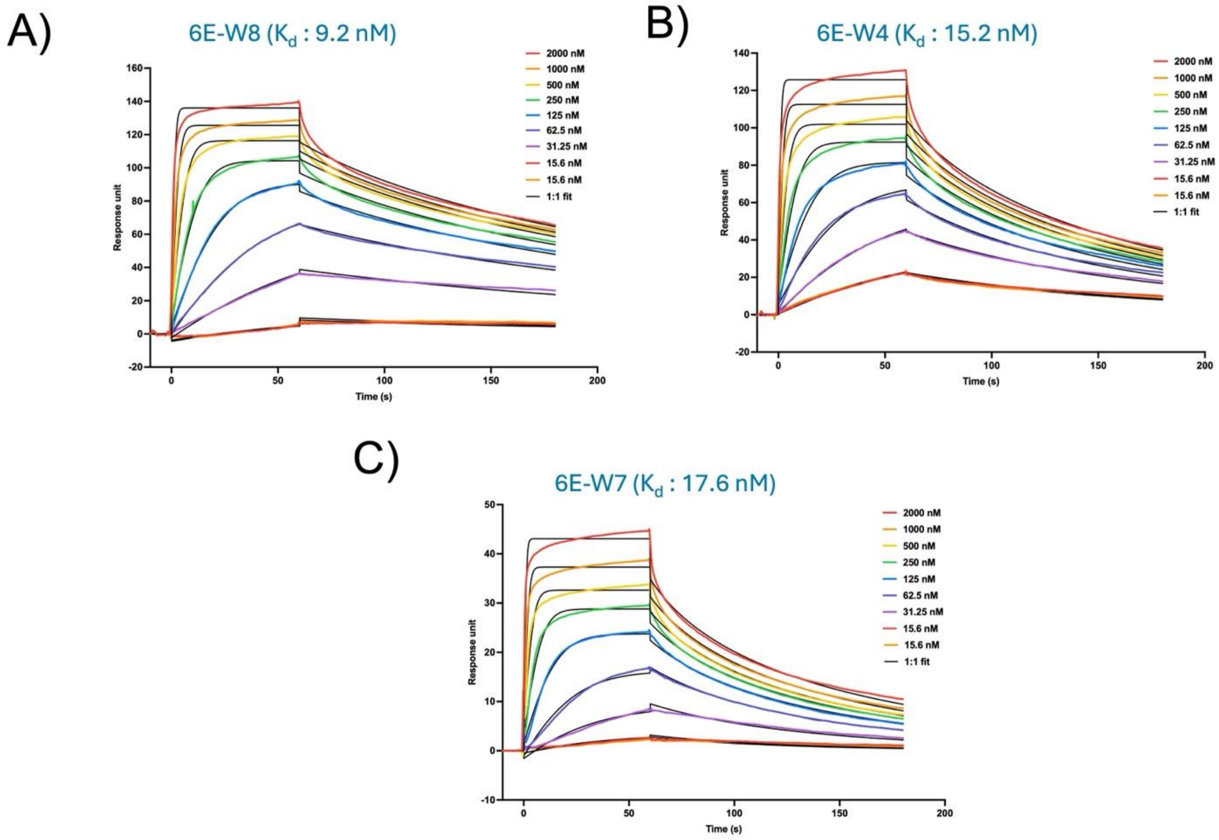


**Supporting Figure 10. Quantitation of binding affinity for 6E variant peptides using SPR.** Representative surface plasmon resonance sensorgrams of 6E variant peptides binding to immobilized $Nb_{6E}$-biotin on a streptavidin chip (Cytiva, #29104992). Colored lines correspond to binding sensorgrams at various peptide concentrations and black lines show the curve-fit generated by local fittings using 1:1 kinetics binding model, which was used to calculate $K_d$ values. Experimental details are described in Methods. Ligand concentrations assessed are as follows: **A)** 6E-W8 (2 - 0.02 μM); **B)** 6E-W4 (2 - 0.02 μM); **C)** 6E-W7 (2 - 0.02 μM).

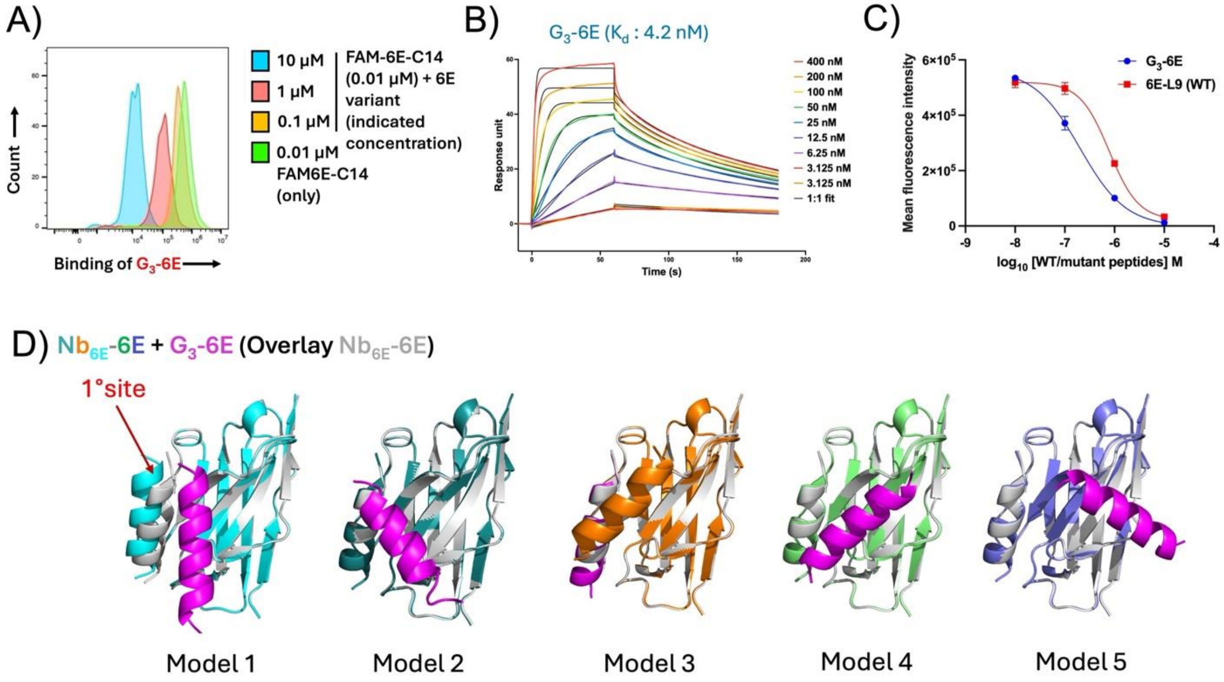


**Supporting Figure 11. Assessment of G$_3$-6E peptide binding and AF modeling. A)** Flow cytometry analysis of G$_3$-6E binding to cells stably expressing A2AR(Nb$_{6E}$) via competition with FAM-6E-C14. Cells were co-incubated with variable concentrations of G$_3$-6E (10–0.01 μM) and a fixed concentration (10 nM) of FAM-6E-C14, followed by washing, and detection with anti-fluorescein antibody conjugated with AF647 (see **Methods**). Histograms depict the intensity of AF647 signaling on live cells from a single representative experiment. **B)** Representative SPR sensorgram of G$_3$-6E (0.4 - 0.003 μM) binding to immobilized Nb$_{6E}$-biotin on a streptavidin coated chip (Cytiva, #29104992). Colored lines correspond to binding sensorgram at various ligand concentrations and black lines denote the curve-fit generated by local fittings using 1:1 kinetics binding model. Experimental details are described in **Methods**. **C)** Experimental flow cytometry competition binding assays to compare 6E-L9 (WT) and G$_3$-6E binding. HEK293 cells stably expressing a cell surface protein fused with Nb$_{6E}$ (A2AR-Nb6E-ALFA) were co-incubated with variable concentrations of 6E-L9 (WT) and G$_3$-6E (10 – 0.01 μM) and a fixed concentration (10 nM) of fluorescein-labelled 6E peptide (FAM-6E-C14), followed by washing, and detection with anti-fluorescein antibody conjugated with Alexafluor647 (AF647) (see **Methods**). Binding was quantified as median AF647 fluorescence intensity. Note that the data for 6E-L9 (WT) is reproduced from main **Figure 5c**. **D**) Evaluation of binding for 6E mutants via competition-based complex formation using AF2 models. Individual AF2 generated models were aligned and overlayed with a control model where 6E peptide occupies the primary (1°) binding site on the Nb$_{6E}$ surface (in grey). G$_3$-6E (magenta) fails to outcompete 6E (in multicolor) for the 1° binding site on Nb$_{6E}$ in four out five models.

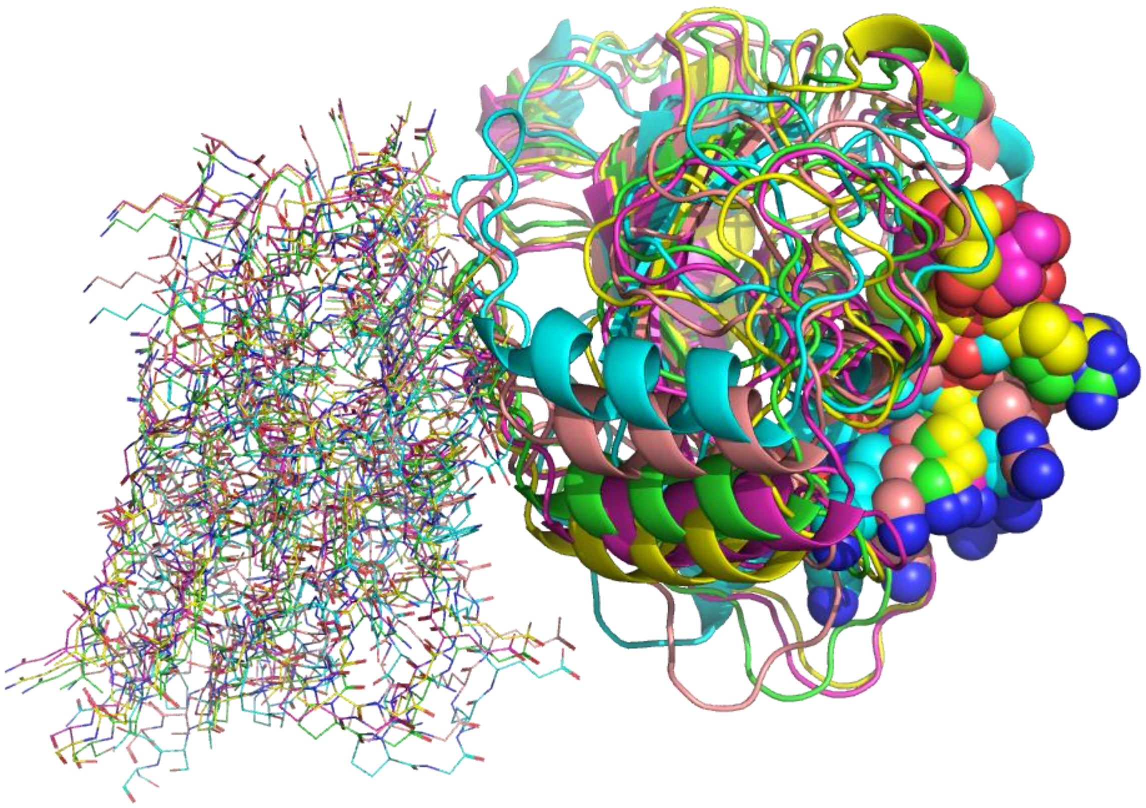

**Supporting Figure 12. Visualization of AF2 models of $Nb_{6E}$-$Ubc_{6E}$ complexes.** Each of 5 AF2 models of is shown in a different color. For each complex $Nb_{6E}$ is found at left shown as sticks, $Ubc_{6E}$ in same color in the shown as cartoon, and the 6E tag in spheres at right. Models were generated by inputting sequences for $Nb_{6E}$ and Ubc6E(1-184), which corresponds to the end of the 6E tag. Residues located C-terminally of the 6E tag are disordered were not included in modeling inputs.

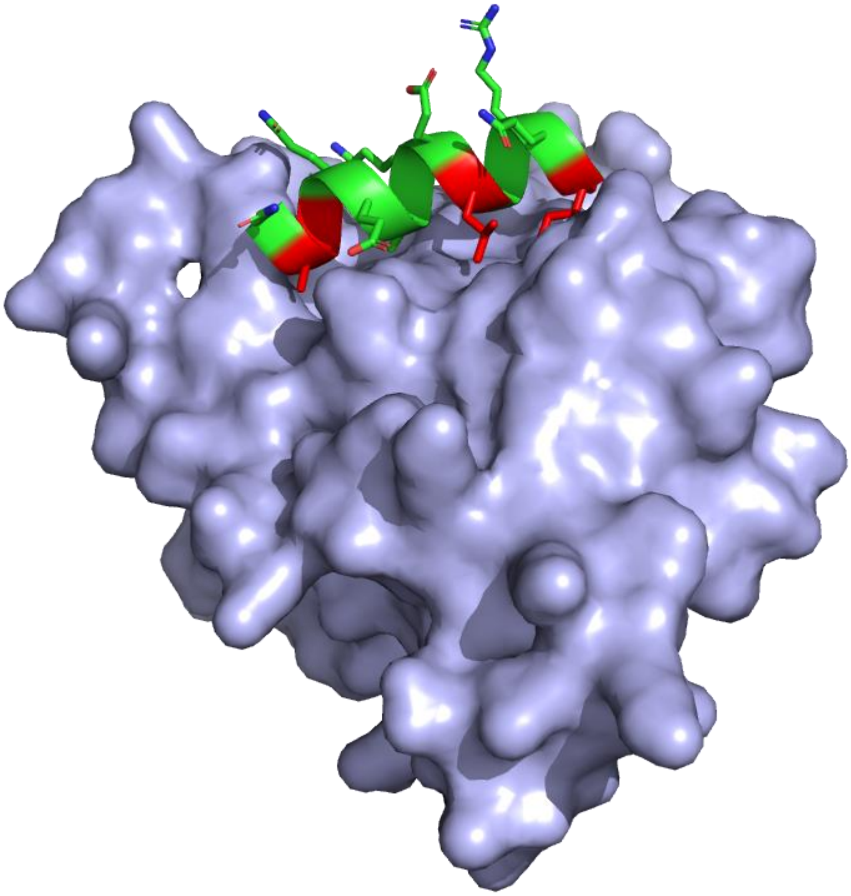

**Supporting Figure 13. Illustration of burial of 6E sidechains in folded Ubc6E structure.** An AF2 model of UBC6E structure is shown. UBC6E is shown as a blue surface except for the tag corresponding to the 6E tag portion, which is shown in green with residues important for binding to Nb6E highlighted in red.
